# Convergent community assembly among globally separated acidic cave biofilms

**DOI:** 10.1101/2022.09.13.507874

**Authors:** Daniel Jones, Irene Schaperdoth, Diana E. Northup, Rodolfo Gómez-Cruz, Jennifer L. Macalady

## Abstract

Acidophilic bacteria and archaea inhabit extreme geochemical ‘islands’ that can tell us when and how geographic barriers affect the biogeography of microorganisms. Here we describe microbial communities from extremely acidic (pH 0-1) biofilms known as “snottites” from hydrogen sulfide-rich caves around the world. Given the extreme acidity and subsurface location of these biofilms, and in light of earlier work showing strong geographic patterns among snottite *Acidithiobacillus* populations, we investigated their structure and diversity in order to understand how geography might impact community assembly. We used 16S rRNA gene cloning and fluorescence *in situ* hybridization (FISH) to investigate 26 snottite samples from four sulfidic caves in Italy and Mexico. All samples had very low biodiversity and were dominated by sulfur-oxidizing bacteria in the genus *Acidithiobacillus. Ferroplasma* and other archaea in the *Thermoplasmatales* ranged from 0 to 50% of total cells, and relatives of the bacterial genera *Acidimicrobium* and *Ferrimicrobium* were up to 15% of total cells. Rare phylotypes included *Sulfobacillus* spp. and members of the *Dependentiae* and *Saccharibacteria* (formerly TM6 and TM7). Although the same genera of acidophiles occurred in snottites on separate continents, most members of those genera represent substantially divergent populations with 16S rRNA genes that are only 95-98% similar. Our findings are consistent with a model of community assembly where sulfidic caves are stochastically colonized by microorganisms from local sources, which are strongly filtered through selection for extreme acid tolerance, and these different colonization histories are maintained by dispersal restrictions within and among caves.

**Importance:** Microorganisms that are adapted to extremely acidic conditions, known as extreme acidophiles, are catalysts for rock weathering, metal cycling, and mineral formation in naturally acidic environments. They are also important drivers of large-scale industrial processes such as biomining and contaminant remediation. Understanding the factors that govern their ecology and distribution can help us better predict and utilize their activities in natural and engineered systems. However, extremely acidic habitats are unusual in that they are almost always isolated within circumneutral landscapes. So where did their acid-adapted inhabitants come from, and how do new colonists arrive and become established? In this study, we took advantage of a unique natural experiment in Earth’s subsurface to show how isolation may have played a role in the colonization history, community assembly, and diversity of highly acidic microbial biofilms.

## Introduction

Extreme acidophiles drive important biogeochemical processes in natural and engineered systems. Acidophilic sulfur- and iron-oxidizing bacteria and archaea speed up the weathering of sulfide minerals in natural ores and mine waste (1, 2), control metal cycling in acidic drainages (3-5), and create secondary mineral deposits in surface and subsurface environments (6, 7). Acidophilic consortia are used for large-scale bioremediation of acidic mine drainage and other heavy metal contaminated environments (8-11), and for industrial processes such as biomining and sulfuric acid production (12-14). Understanding the ecology of organisms in naturally acidic environments can inform microbial behavior in engineered systems and provide ideas for new bioremediation strategies (e.g., 11, 15). Furthermore, acidic environments are useful model systems for studying microbial evolution and ecology because they often contain low biodiversity (16) and represent extreme geochemical “islands” in a sea of neutrophilic habitats (17).

Studies conducted at local scales in acidic systems have identified geochemical factors that influence microbial distribution, including pH, conductivity, and metal concentrations (e.g., 18-25). However, microbial spatial distribution might not only be a result of natural selection by contemporary environmental factors, but also a legacy of past environmental conditions and stochastic colonization events (26). Dispersal barriers limit the movement of new species into an environment, and as a result, communities from the same geographic region are more likely to contain the same organisms because they share a similar biotic history (27-29).

Extremely acidic microbial biofilms form on the walls and ceilings of sulfide-rich caves, in areas where hydrogen sulfide degasses from springs and streams to the cave atmosphere (Figure 1). These biofilms, commonly known as “snottites,” have pH values between 0 and 1.5, and are found hanging from the surface of gypsum corrosion residues that isolate them from buffering by the parent limestone. Snottites have been consistently observed where H_2_S(*g*) concentrations are between 0.2 to 25 parts-per-million by volume (ppmv), and rarely occur where concentrations are outside of this range (30, 31). Previous research has shown that snottites are inhabited by very low biodiversity communities that contain abundant *Acidithiobacillus* spp., as well as representatives or close relatives of the genera *Acidimicrobium, Sulfobacillus, Ferroplasma*, and *Cuniculiplasma* (30-35).

**Figure 1.**
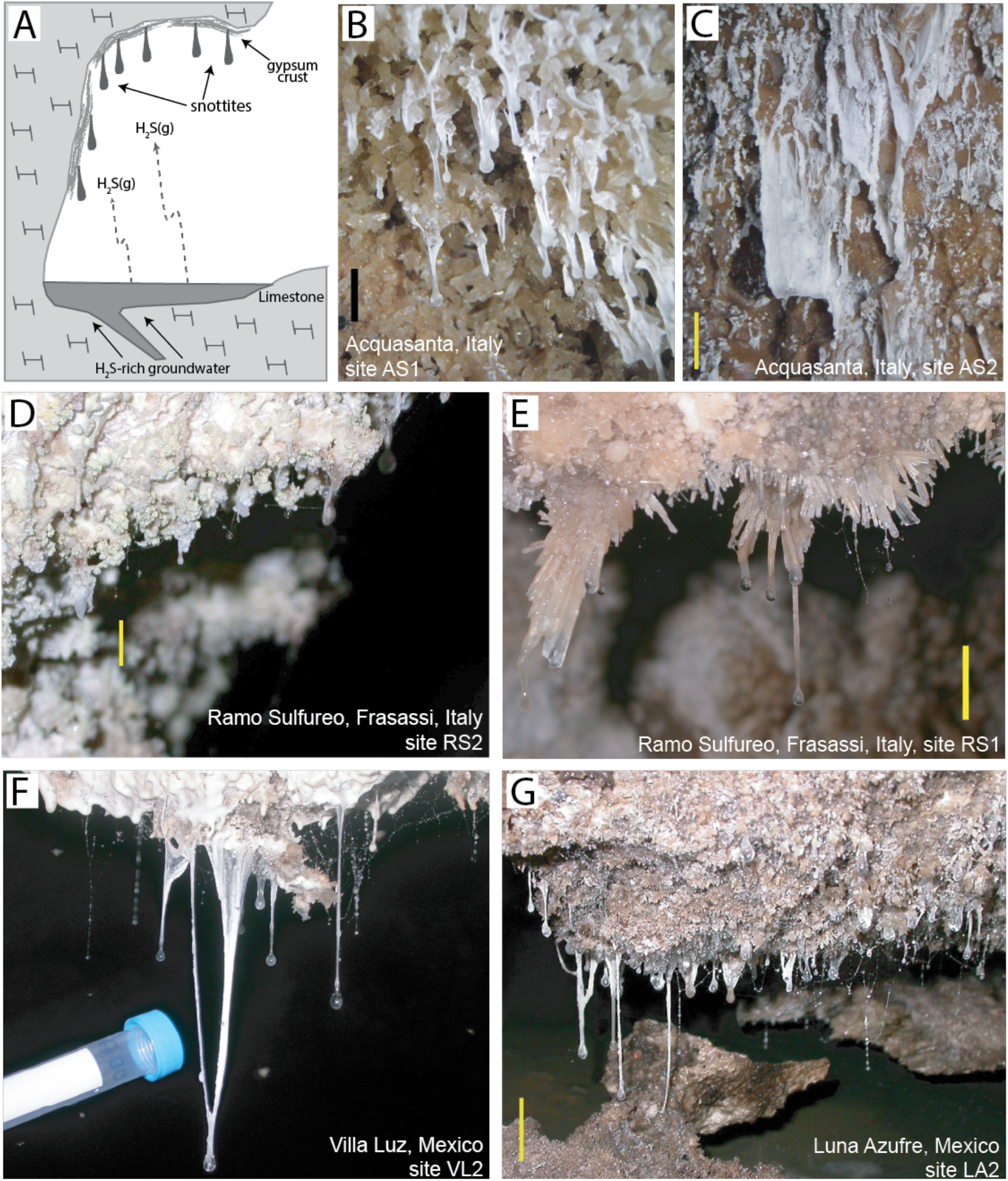
(A) Schematic depicting the snottite niche in sulfidic caves. Snottites hang from gypsum wall crusts on cave walls and ceilings in close proximity to sulfidic cave streams. (B-G) Representative photographs of snottite communities described in this study. Yellow scale bars are 2 cm, black bar in (b) is 1 cm, and the collection tube in (f) is 1.5 cm in diameter.

Sulfidic cave snottites present an exceptional opportunity to test how geographic barriers affect microbial communities. Because snottites occur in stable subsurface environments and only form on overhanging surfaces within a specific range of H_2_S(*g*) concentrations, geochemical variability among snottite locations is relatively limited (Figure 1a). Furthermore, their extremely low pH is a potential barrier to microbial dispersal in addition to the physical barrier imposed by their subsurface location. In previous research, Jones et al. (36) used high-resolution genetic analyses to resolve relationships among geographically separated populations of *Acidithiobacillus spp*. from cave snottites. That work showed that *Acidithiobacillus* populations from separate cave locations are divergent, and identified a distance-decay relationship among the most abundant *Acidithiobacillus* spp. that was consistent with restricted dispersal among cave locations. We therefore hypothesized that geographic isolation would also affect other organisms in the snottite biofilms, and thereby impact the structure and diversity of the whole biofilm community. We tested this hypothesis by sampling snottites from four caves on two continents, and quantitatively comparing community structure and composition using 16S rRNA gene sequencing and fluorescence *in situ* hybridization (FISH).

## Results

### Field observations and geochemistry

Snottites were collected from multiple locations in four sulfidic caves: Cueva de Villa Luz and Cueva Luna Azufre in Tabasco, Mexico, and the Grotte del Fiume (Frasassi) and Grotta Nuova di Rio Garrafo (Acquasanta Terme) in the Marche region, Italy. H_2_S(*g*) concentrations at sampling sites ranged from 0.2 to 24 ppmv (Table 1), except for samples GS07-35 and AS08-5 where H_2_S(*g*) was below detection. Snottites were also observed (but not collected) in the Yellow Roses room of Cueva de Villa Luz, at H_2_S(*g*) concentrations between 30-50 ppm. H_2_S(*g*) commonly exceeds 80 ppm in the atmosphere of this enclosed room, and snottites are not observed under these conditions (30, 35). Similarly, no snottites were observed near a thermal cave stream in Acquasanta, where H_2_S(*g*) concentrations commonly exceed 100 ppm (37, 38). We have observed that snottites form on a time scale as short as months to weeks in Frasassi and Acquasanta, and at Villa Luz they have been observed to form in days (39).

**Table 1.** Summary of snottite samples and associated environmental data.

| Sample | Cave | Site | Wall mineralogy | Temp (°C) | [H<sub>2</sub>S(g)] (ppmv) | [CO<sub>2</sub>(g)] | [SO<sub>2</sub>(g)] (ppmv) |
| --- | --- | --- | --- | --- | --- | --- | --- |
| GS07-35 | Frasassi | GS2 | gypsum | 13-14 | bd<sup>b</sup> | 700 ppm | bd |
| GS08-6 | Frasassi | GS1 | gypsum | 13-14 | 0.8 (0.6-1) | 1400 ppm | nm |
| RS05-2 | Frasassi | RS2 | gypsum and S<sup>o</sup> | 13-14 | nm | nm | nm |
| RS05-24 | Frasassi | RS2 | gypsum and S<sup>o</sup> | 13-14 | 16 (8-24) | >3000 ppm | nm |
| RS05-1 | Frasassi | RS1 | gypsum | 13-14 | nm | nm | nm |
| RS05-20 | Frasassi | RS1 | gypsum | 13-14 | 3.4 | >3000 ppm | nm |
| RS07-21 | Frasassi | RS3 | gypsum and S<sup>o</sup> | 13-14 | 21 (17-25) | 0.2 % | bd |
| RS07-22 | Frasassi | RS2 | gypsum and S<sup>o</sup> | 13-14 | 14.5 (11-18) | 0.2 % | bd |
| RS08-30 | Frasassi | RS2 | gypsum and S<sup>o</sup> | 13-14 | 11.5 (11-12) | 2500 ppm | nm |
| RS08-31 | Frasassi | RS3 | gypsum and S<sup>o</sup> | 13-14 | 14 (12-16) | 3000 ppm | nm |
| RS09-1 | Frasassi | RS2 | gypsum and S<sup>o</sup> | 13-14 | 8 (5-11) | 0.25% | bd |
| RS09-10 | Frasassi | RS2 | gypsum and S<sup>o</sup> | 13-14 | 10.5 (8-13) | 0.3% | bd |
| RS09-11 | Frasassi | RS3 | gypsum and S<sup>o</sup> | 13-14 | 21 (20-22) | 0.4% | bd |
| PC05-1 | Frasassi | PC1 | gypsum | 13-14 | nm | nm | nm |
| PC05-13 | Frasassi | PC1 | gypsum | 13-14 | 1.1 (0.2-2) | 1200 ppm | nm |
| GB08-30 | Frasassi | GB1 | gypsum | 13-14 | 4 (3-5) | 1400 ppm | nm |
| AS05-5 | Acquasanta | AS3 | gypsum | 19-21 | nm | nm | nm |
| AS07-1 | Acquasanta | AS3 | gypsum | 19-21 | 2 (1-3) | 0.4 % | 4.5 (2-36) |
| AS07-3 | Acquasanta | AS4 | sediment<sup>a</sup> | 19-21 | 2.9 (1-4.8) | 0.4 % | 5 (3-7) |
| AS07-4 | Acquasanta | AS2 | sediment<sup>a</sup> | 25-35 | 24 (18-30) | 0.6 % | 11.5 (10-13) |
| AS08-5 | Acquasanta | AS3 | gypsum | 19-21 | bd<sup>c</sup> | 0.45 % | 5 (4-6) |
| AS08-6 | Acquasanta | AS2 | sediment<sup>a</sup> | 25-35 | 3 | 0.55 % | 4.5 (4-5) |
| VL07-20 | Villa Luz | VL1 | gypsum | 25-28 | 12 (6-18) | 1200 ppm | bd |
| VL07-26 | Villa Luz | VL2 | gypsum | 25-28 | 3.5 (3-4) | 1100 ppm | bd |
| LA07-10 | Luna Azufre | LA1 | gypsum | 28-33 | 3 | 1250 ppm | bd |
| LA07-13 | Luna Azufre | LA2 | gypsum | 28-33 | 2 | 1250 ppm | bd |

<sup>a</sup>Sediment is a mixture of gypsum, clay, and chert particles
<sup>b</sup>Measured by ENMET MX2100, lower detection limit (ldl) 1 ppm.
<sup>c</sup>Measured by Draeger tube, ldl 0.2 ppm.

### 16S rRNA gene libraries

We cloned archaeal and bacterial 16S rRNA genes from six snottite samples (Table S1). *Acidithiobacillus* phylotypes (Figure 2) were the most abundant clones in all bacterial libraries, and were the only bacterial sequences retrieved from samples PC05-1, AS07-3, and AS08-5. The other three libraries contained relatives of the bacterial genera *Acidimicrobium* and *Ferrimicrobium* (Figure 3). Libraries from samples RS05-2 and VL07-20 also contained phylotypes of the genus *Sulfobacillus* and candidate phylum *Dependentiae* (formerly TM6 (40)) (Figure S1, Figure S2). Samples VL07-20 and LA07-10 also contained members of the *Saccharibacteria* (formerly candidate phylum TM7 (41)). *Ferroplasma* relatives are the most abundant archaeal clones in universal and archaeal-specific clone libraries (Figure 4). Other archaeal clones are related to “C-plasma” and *Cuniculiplasma* (formerly “G-plasma” (3, 42)) in the order *Thermoplasmatales* (Figure 4).

**Figure 2.**
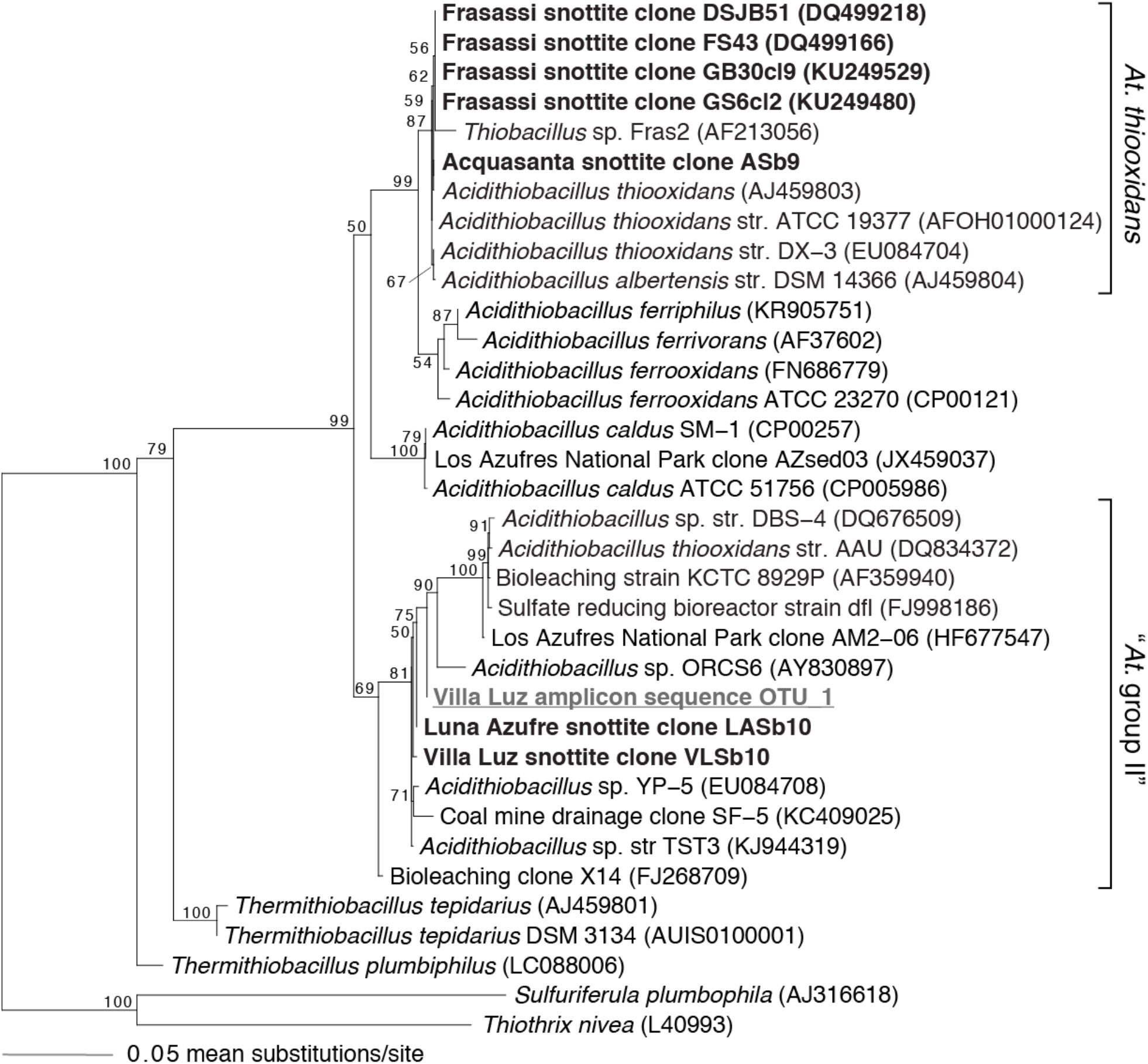
Phylogenetic analysis of 16S rRNA gene sequences from the genus *Acidithiobacillus*. The base tree is a maximum likelihood phylogram created with nearly full-length sequences, with shorter amplicon sequence (gray underline) placed using the EPA algorithm. Representative sequences from 16S rRNA gene clones are in bold black. Bootstrap values >50% are provided for each node.

**Figure 3.**
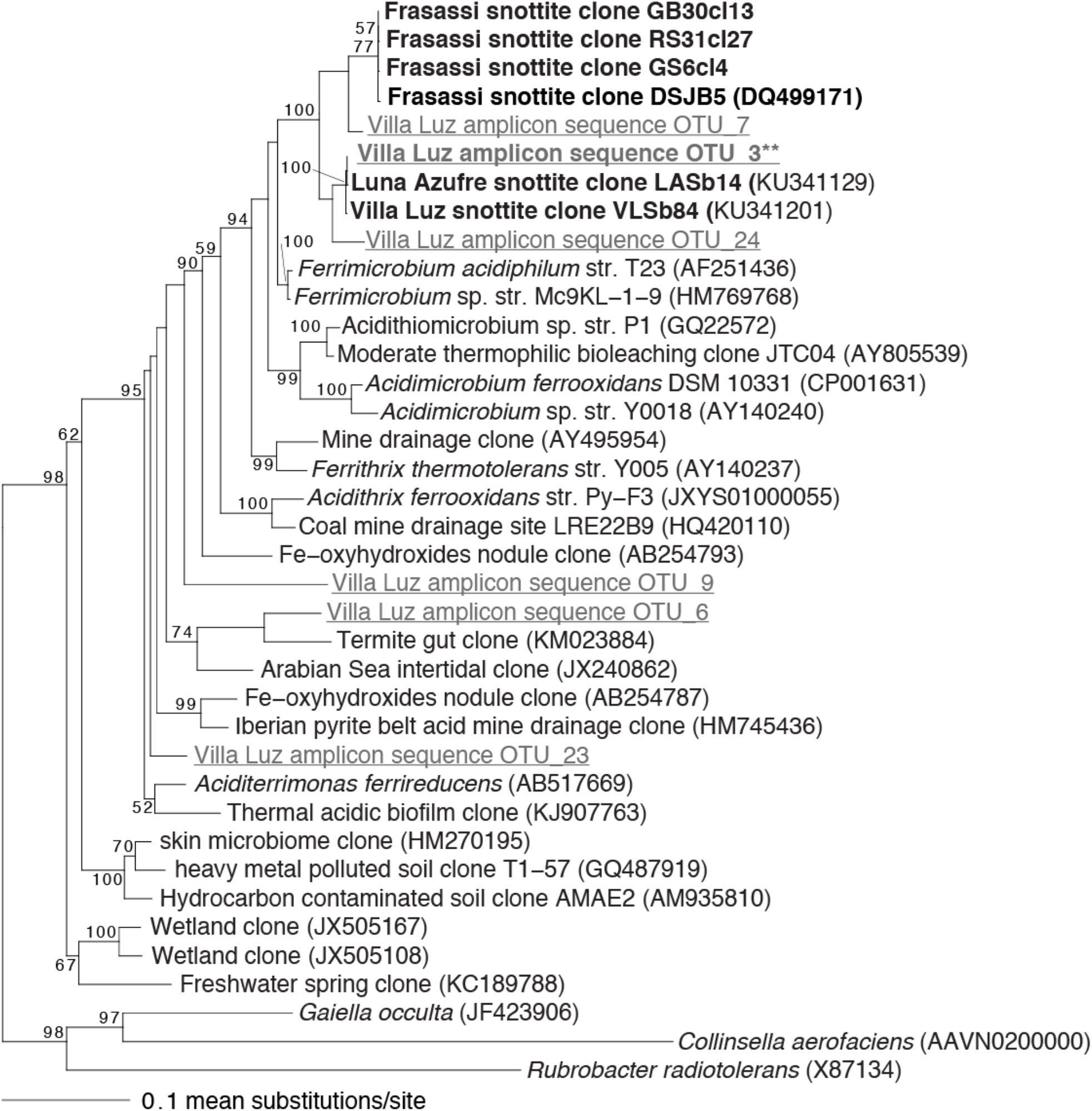
Phylogenetic analysis of 16S rRNA gene sequences from the *Acidimicrobiaceae* family. The base tree is a maximum likelihood phylogram created with nearly full-length sequences, with shorter amplicon sequences (gray underline) placed after the fact using the EPA algorithm. Representative sequences from 16S rRNA gene clones are in bold black. The amplicon sequence in bold gray indicated by two asterisks (**) represents that most abundant *Acidimicrobiaceae* OTU (24.8 and 13.7% of the VL13-1 and VL13-2 libraries, respectively); other *Acidimicrobiaceae* OTUs are <2% relative abundance (Table S2). Bootstrap values >50% are provided for each node.

**Figure 4.**
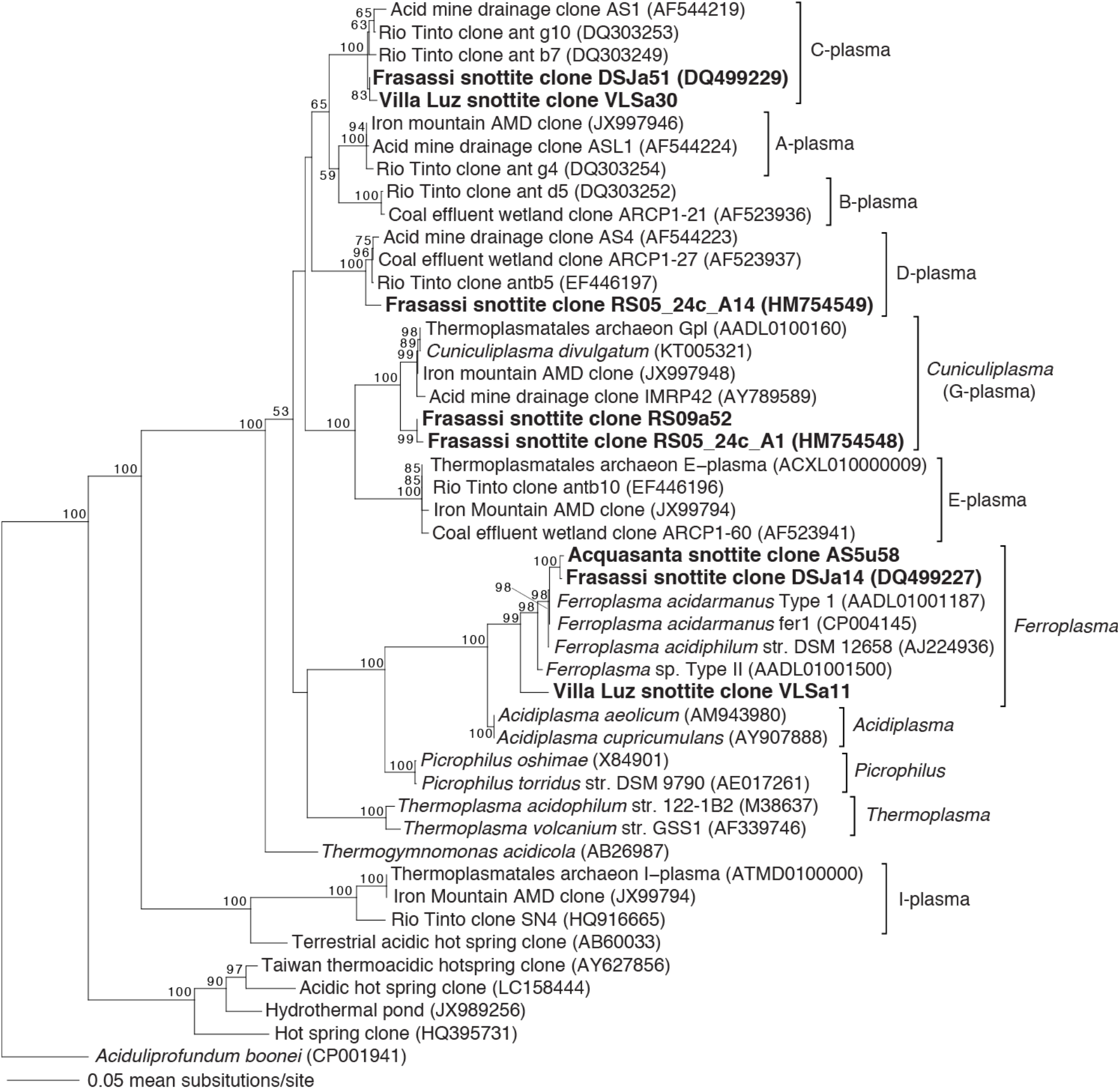
Neighbor joining phylogram of 16S rRNA gene sequences from the Archaeal family *Thermoplasmatales*. Snottite clones are in bold black. Bootstrap values >50% are provided for each node. Group names for the ‘alphabet plasmas’ are after Baker and Banfield (3).

Although many of the same microbial genera were present in snottites from both Italy and Mexico, snottite clones from the two countries have less than 97% 16S rRNA gene sequence identity (Table S1). *Acidithiobacillus* clones are <95% similar (Figure 2), clones affiliated with *Acidimicrobium/Ferrimicrobium* are <95% similar (Figure 3), *Sulfobacillus* clones are <92% similar (Figure S1), and *Ferroplasma* sequences are <97% similar (Figure 4). The only exception are C-plasma clones VLSa30 and DSJa51, which share 99.6% similarity (Figure 3, Table S1), and clone DSJb30 in the *Dependentiae*, which is 97.9-98.1% similar to Villa Luz clones VLSb7, VLSb29, and VLSb87 (Figure S2).

All of the *Acidithiobacillus* clones from Italy are members of the genus *Acidithiobacillus thiooxidans*, while all *Acidithiobacillus* clones from the Mexican caves represent a separate species of *Acidithiobacillus* (referred to as “*At*. group II“ in Figure 2, after Jones et al. (2016)) that is more closely related to *At. caldus* than *At. thiooxidans*. In order to see if deeper sequencing would reveal the presence of *At. thiooxidans* in Cueva de Villa Luz, we generated much larger rRNA gene libraries from samples collected during a later expedition to Villa Luz. *At. thiooxidans* sequences were not detected among 143,944 and 102,229 rRNA gene amplicon sequences from two snottite samples. Like the clone sequences, *Acidithiobacillus* spp. in these samples formed a single operational taxonomic unit (OTU) that was similar to clone sequences in *At*. group II. Other abundant OTUs in the deeper amplicon datasets include members of groups that were identified in clone libraries (*Acidimicrobiaceae, Sulfobacillus, Saccharibacteria, Dependentiae*), as well as other members of the *Gamma*- and *Alphaproteobacteria* that were not identified in the clones libraries, including an abundant OTU of *Metallibacterium* and multiple OTUs from the *Acidimicrobiaceae* (Figure 2, 3, S1, S2, Table S2).

### Fluorescence *in situ* hybridization (FISH)

We sampled 26 snottite communities from 14 cave locations using FISH (Table 2). Representative FISH photomicrographs are shown in Figures 5 and 6. All samples contain over 50% THIO1-positive cells (*Acidithiobacillus*), with the exception of AS05-5. In sample AS05-5, over 25% of DAPI-strained cells did not hybridize with any FISH probes, possibly indicating that a large proportion of the community was inactive at the time of collection. ACM732-positive cells (*Acidimicrobium*/*Ferrimicrobium*) are present in most but not all samples, and make up to 13.7% of total cells. In most samples, bacterial cells that do not hybridize with either THIO1 or ACM732, hereafter referred to as “other bacteria,” are <3% of cells. However, samples RS08-31, LA07-10, and VL07-26 have between 5.9-6.9% other bacteria, and sample VL07-20 contains 15% other bacteria. At least some archaea are present in all samples, although some samples contain less than 2% ARCH915-positive cells. Twelve samples contain >2% FER656-positive cells (*Ferroplasma*), including four Acquasanta samples with between 25-49% FER656. However, *Ferroplasma* are absent from some samples that have abundant ARCH915-positive cells. Non-*Ferroplasma* archaea make up 15-30% of total cells in several samples, including AS08-6 and several Frasassi RS samples.

**Table 2.**
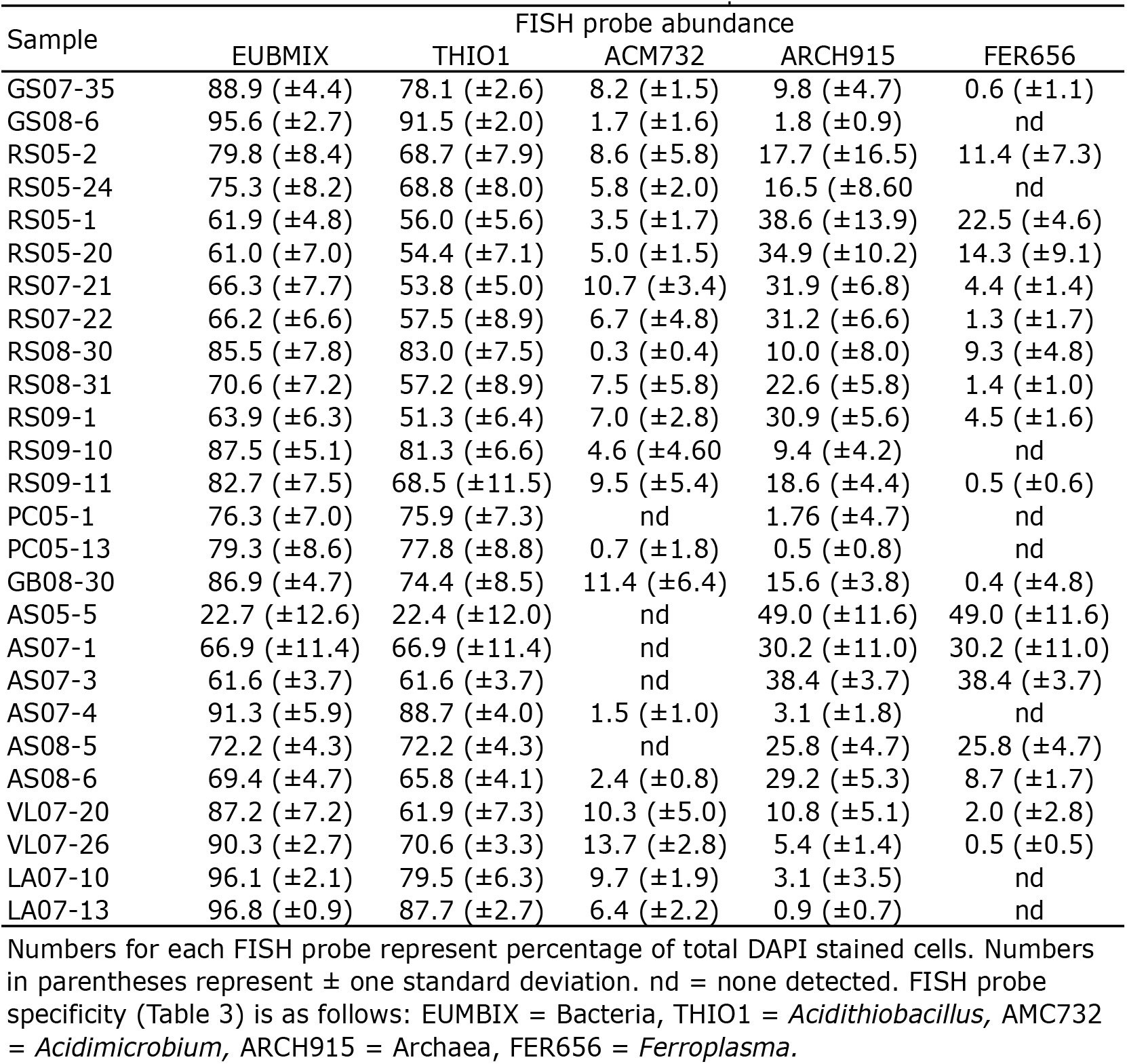
FISH cell counts for snottite communities

| Sample | FISH probe abundance |  |  |  |  |
| --- | --- | --- | --- | --- | --- |
|  | EUBMIX | THIO1 | ACM732 | ARCH915 | FER656 |
| GS07-35 | 88.9 (±4.4) | 78.1 (±2.6) | 8.2 (±1.5) | 9.8 (±4.7) | 0.6 (±1.1) |
| GS08-6 | 95.6 (±2.7) | 91.5 (±2.0) | 1.7 (±1.6) | 1.8 (±0.9) | nd |
| RS05-2 | 79.8 (±8.4) | 68.7 (±7.9) | 8.6 (±5.8) | 17.7 (±16.5) | 11.4 (±7.3) |
| RS05-24 | 75.3 (±8.2) | 68.8 (±8.0) | 5.8 (±2.0) | 16.5 (±8.60) | nd |
| RS05-1 | 61.9 (±4.8) | 56.0 (±5.6) | 3.5 (±1.7) | 38.6 (±13.9) | 22.5 (±4.6) |
| RS05-20 | 61.0 (±7.0) | 54.4 (±7.1) | 5.0 (±1.5) | 34.9 (±10.2) | 14.3 (±9.1) |
| RS07-21 | 66.3 (±7.7) | 53.8 (±5.0) | 10.7 (±3.4) | 31.9 (±6.8) | 4.4 (±1.4) |
| RS07-22 | 66.2 (±6.6) | 57.5 (±8.9) | 6.7 (±4.8) | 31.2 (±6.6) | 1.3 (±1.7) |
| RS08-30 | 85.5 (±7.8) | 83.0 (±7.5) | 0.3 (±0.4) | 10.0 (±8.0) | 9.3 (±4.8) |
| RS08-31 | 70.6 (±7.2) | 57.2 (±8.9) | 7.5 (±5.8) | 22.6 (±5.8) | 1.4 (±1.0) |
| RS09-1 | 63.9 (±6.3) | 51.3 (±6.4) | 7.0 (±2.8) | 30.9 (±5.6) | 4.5 (±1.6) |
| RS09-10 | 87.5 (±5.1) | 81.3 (±6.6) | 4.6 (±4.60) | 9.4 (±4.2) | nd |
| RS09-11 | 82.7 (±7.5) | 68.5 (±11.5) | 9.5 (±5.4) | 18.6 (±4.4) | 0.5 (±0.6) |
| PC05-1 | 76.3 (±7.0) | 75.9 (±7.3) | nd | 1.76 (±4.7) | nd |
| PC05-13 | 79.3 (±8.6) | 77.8 (±8.8) | 0.7 (±1.8) | 0.5 (±0.8) | nd |
| GB08-30 | 86.9 (±4.7) | 74.4 (±8.5) | 11.4 (±6.4) | 15.6 (±3.8) | 0.4 (±4.8) |
| AS05-5 | 22.7 (±12.6) | 22.4 (±12.0) | nd | 49.0 (±11.6) | 49.0 (±11.6) |
| AS07-1 | 66.9 (±11.4) | 66.9 (±11.4) | nd | 30.2 (±11.0) | 30.2 (±11.0) |
| AS07-3 | 61.6 (±3.7) | 61.6 (±3.7) | nd | 38.4 (±3.7) | 38.4 (±3.7) |
| AS07-4 | 91.3 (±5.9) | 88.7 (±4.0) | 1.5 (±1.0) | 3.1 (±1.8) | nd |
| AS08-5 | 72.2 (±4.3) | 72.2 (±4.3) | nd | 25.8 (±4.7) | 25.8 (±4.7) |
| AS08-6 | 69.4 (±4.7) | 65.8 (±4.1) | 2.4 (±0.8) | 29.2 (±5.3) | 8.7 (±1.7) |
| VL07-20 | 87.2 (±7.2) | 61.9 (±7.3) | 10.3 (±5.0) | 10.8 (±5.1) | 2.0 (±2.8) |
| VL07-26 | 90.3 (±2.7) | 70.6 (±3.3) | 13.7 (±2.8) | 5.4 (±1.4) | 0.5 (±0.5) |
| LA07-10 | 96.1 (±2.1) | 79.5 (±6.3) | 9.7 (±1.9) | 3.1 (±3.5) | nd |
| LA07-13 | 96.8 (±0.9) | 87.7 (±2.7) | 6.4 (±2.2) | 0.9 (±0.7) | nd |

**Figure 5.**
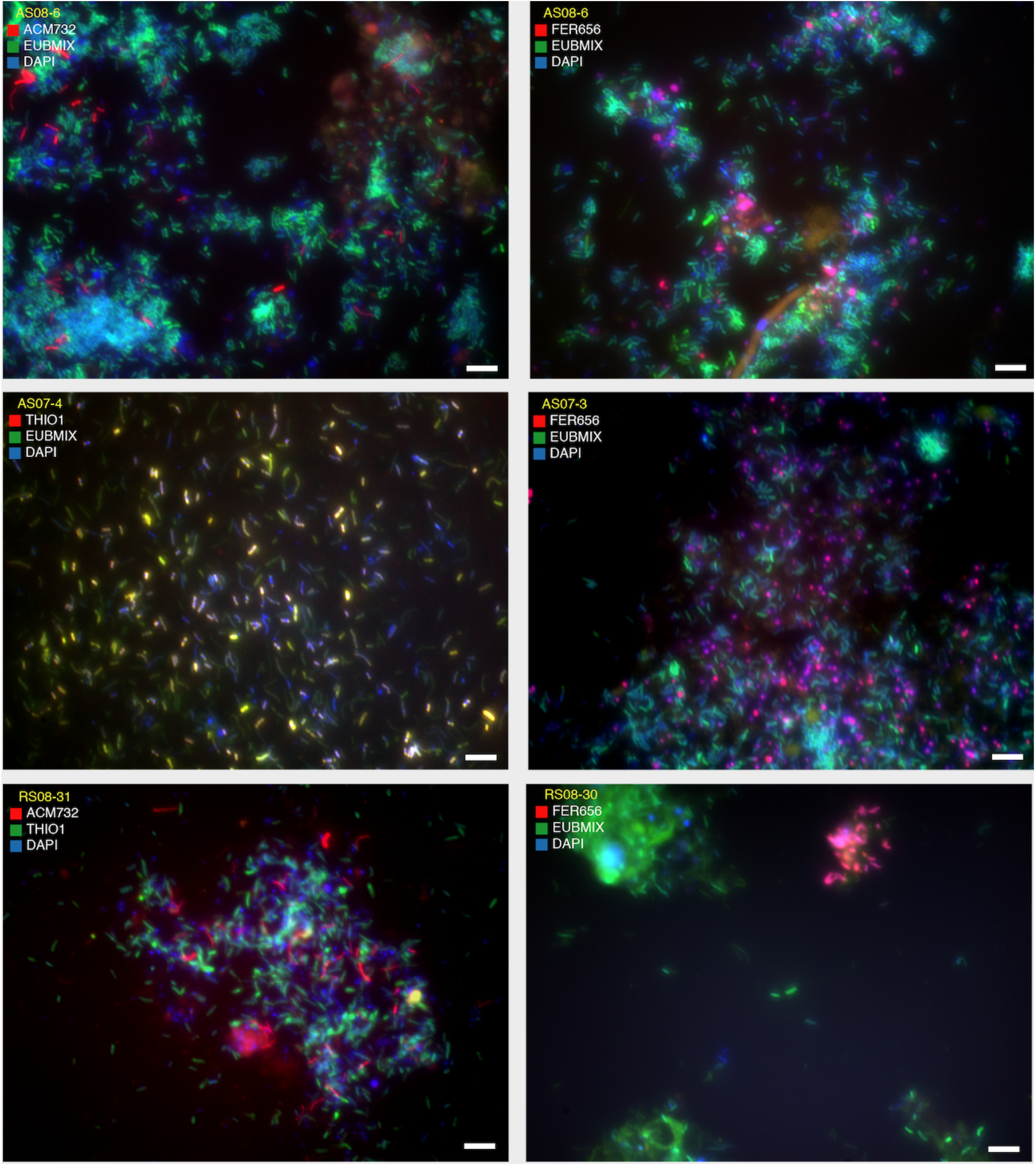
Representative fluorescence *in situ* hybridization (FISH) photomicrographs. The sample name and probes used for each photo are provided in the legend. White scale bars are 5 μM. Abundant DAPI-labeled cells in AS08-6 are nearly all archaea, as are abundant DAPI-labeled cells in RS08-31.

**Figure 6.**
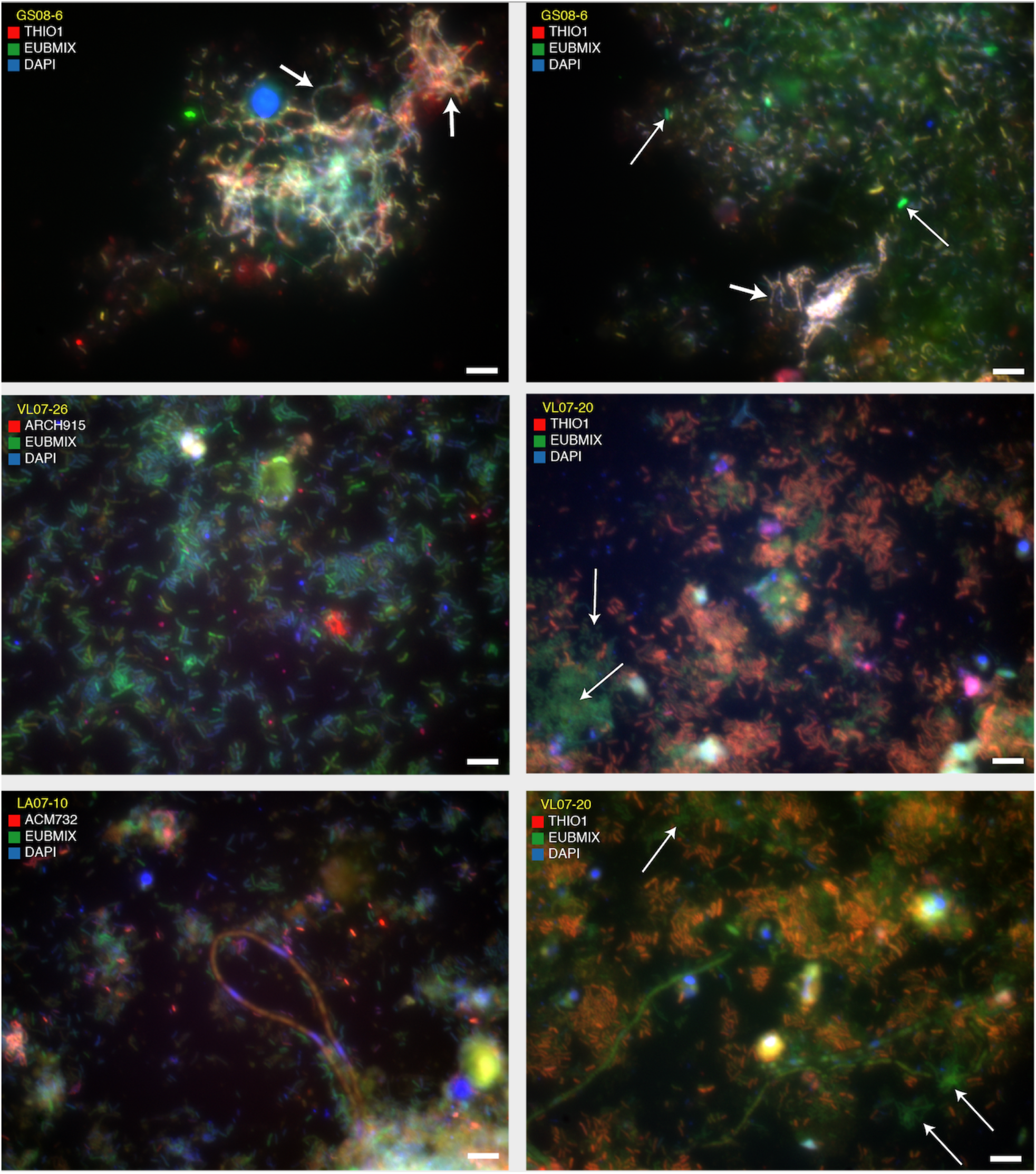
Representative fluorescence *in situ* hybridization (FISH) photomicrographs. The sample name and probes used for each photo are provided in the legend. White scale bars are 5 μM. Note the abundant non-THIO1 labeled cells (white arrows) in VL07-20. Fat arrow in GS08-6 indicates THIO1-labeled cells linked into filaments, and thin arrows indicate non-THIO1-labeled bacterial cells.

We used principal component analysis (PCA) to explore the major axes of variance in community composition based on the FISH dataset (Figure 7). For this and other statistical analyses using FISH data (Table 2), our “species” were cell counts of THIO1, ACM732, FER656, “other bacteria” (EUBMIX – (THIO1 + ACM732)), and “other archaea” (ARCH915 – FER656). Principal components 1 and 2 combined explain more than 75% of the total variance. The first principal component in Figure 7 appears to be related to location, with Acquasanta samples plotting opposite Villa Luz samples. Species loadings show that the taxa driving this clustering are FER656-positive cells, which were most abundant in the Acquasanta samples, and ACM732 and “other bacteria” that were more abundant in Villa Luz.

**Figure 7.**
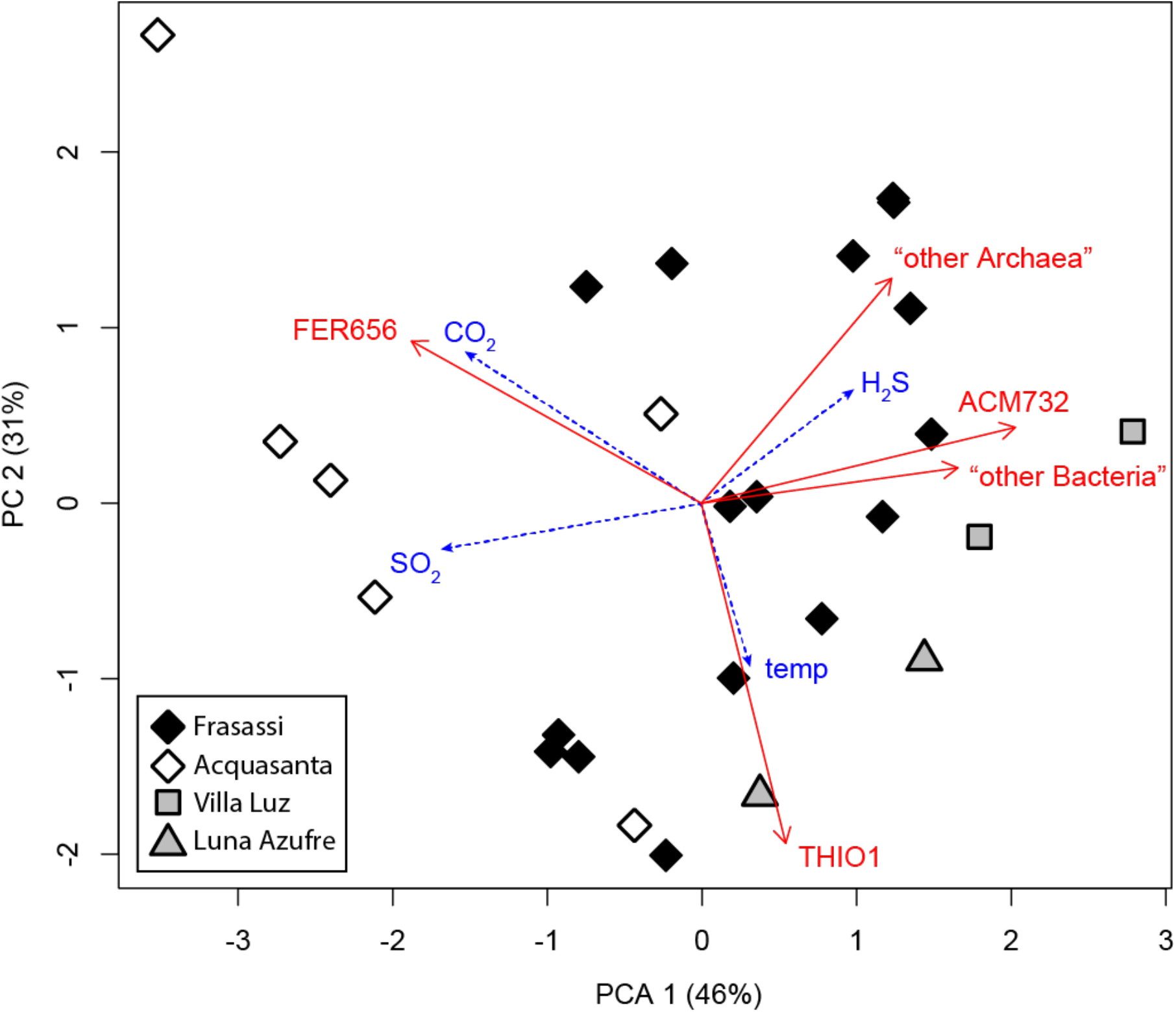
Principal component analysis (PCA) of FISH cell counts (Table 3). Sample scores are coded by cave (communities from different caves are significantly different by ANOSIM). Fitted vectors of environmental variables are included as an overlay; CO_2_ and SO_2_ gas concentrations are statistically significantly correlated with the ordination axes (p <0.05), H_2_S gas and temperature are not (p > 0.1).

ANalysis Of SIMilarities (ANOSIM; McCune and Grace, 2002) of FISH data indicates that communities from different caves are significantly different (ANOSIM R = 0.268, p = 0.013), and this result remains significant if Villa Luz and Luna Azufre samples are pooled (ANOSIM R = 0.287, p = 0.006). Mantel tests show that there is a statistically significant correlation between community dissimilarity and geographical distance, both for all samples and for Italian samples alone, if rank correlation is used (Mantel test with Spearman’s *rho*: all samples, *rho*=0.174, p=0.045; Italian samples only, *rho*=0.358, p<0.001; Frasassi samples only, *rho*=0.158, p=0.065).

**Table 3.**
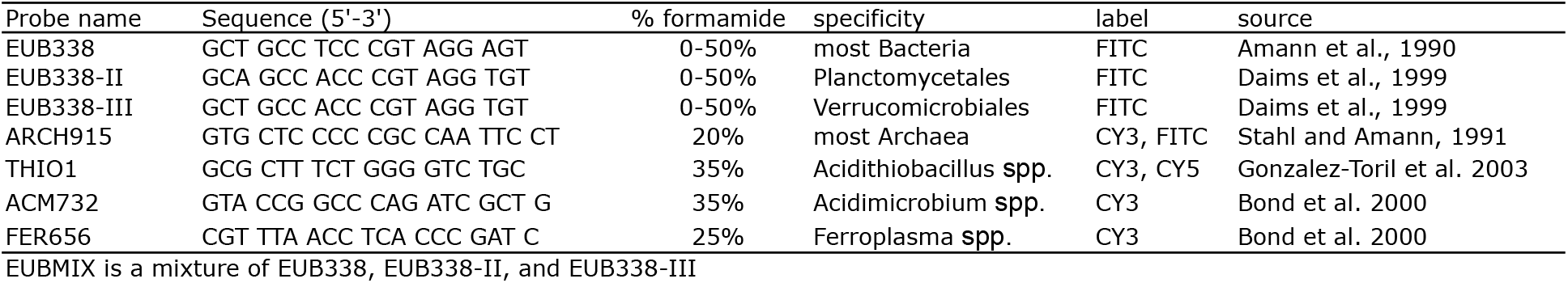
FISH probes used.

We also examined whether community composition is related to the concentration of gases in the cave air. When these variables are fitted to the ordination analysis, CO_2_ and SO_2_ concentration align with the first ordination axis in the direction of the Acquasanta samples (Figure 7), consistent with the high gas concentrations in that cave. These two variables are statistically significantly correlated with the ordination axes (r^2^ = 0.33, p = 0.02 for CO_2_; r^2^ = 0.31, p = 0.02 for SO_2_), while H_2_S concentration and temperature are not (r^2^ = 0.15, p = 0.22 for H_2_S; r^2^ = 0.10, p = 0.34 for temperature).

## Discussion

### Snottite microbial communities

*Acidithiobacillus* spp. were more than 50% of total cells in all except one of the 26 snottite samples analyzed with FISH, and they were frequently more than 80% of total cells (Table 2). *Acidithiobacillus* was also the only group present in all of the biofilms sampled. These findings are consistent with other studies of snottite biofilms from Frasassi and Cueva de Villa Luz (30, 31, 34, 35). Based on previous metagenomic and culture-based analyses, *Acidithiobacillus thiooxidans* in Italian snottites are autotrophic sulfide-oxidizers with the capacity to form extracellular polymeric substances, and were described as possible ‘architects’ of the biofilms (32, 36). The predominance of *Acidithiobacillus* spp. across the large number of samples analyzed here is further evidence that *Acidithiobacillus* are responsible for forming the acidic biofilms, and that they are important organisms for sulfide oxidation and sulfuric acid production in sulfidic caves.

Close relatives or members of the bacterial genera *Acidimicrobium* and *Ferrimicrobium* were present in most samples. Based on metagenomic analyses of Frasassi snottites, these populations do not have the capacity to fix inorganic carbon, and are likely chemoorganoheterotrophs or chemolithoheterotrophs (32, 33). However, the coverage of these populations in previous snottite metagenomes was very low, and 16S rRNA genes recovered from snottite populations are divergent from known isolates within the *Acidimicrobiaceae* family (Figure 2), so inferences about their physiology are necessarily limited.

Other bacterial members included *Sulfobacillus* and members of the candidate phyla *Dependentiae* (formerly TM6) and *Saccharibacteria* (formerly TM7). We hypothesize that, based on their closest relatives in phylogenetic analyses and other members of their genera, the *Sulfobacillus* represent another inorganic sulfur-oxidizing or organoheterotrophic population (e.g., 43, 44) and the *Dependentiae* and *Saccharibacteria* are likely obligate symbionts or parasites (40, 45, 46) (Figure 2; Figure S1). The protists and fungi that are commonly observed in snottites (Figures 5, 6, S3; (31)) could be hosts for symbiotic *Dependentiae* and *Saccharibacteria*. We observed nematodes and arthropods in Villa Luz and Luna Azufre snottites that are additional potential hosts for symbionts (Figure S3). These likely represent the extremely acid-tolerant bacteriophagous nematode (*Mesorhabditis acidophila*) and nematode-eating mites previously described from Villa Luz snottites (47).

Archaea were present in all of the snottites sampled (Table 2). Members of the genus *Ferroplasma* were detected in 18 out of the 26 samples analyzed using FISH, and were up to 50% of total cells. However, we also observed significant numbers of ARCH915-positive cells that did not hybridize with FER656. Other archaeal populations identified in this and previous studies of snottites include ‘C-plasma’ and *Cuniculiplasma* (formerly ‘G-plasma’) in the *Thermoplasmatales* (Figure 4) (32, 33). In previous metagenomic analysis, *Ferroplasma* from the Acquasanta caves had the genetic potential for sulfur oxidation, and *Cuniculiplasma* from Frasassi had a homologue of sulfide quinone oxidoreductase (33), so these archaeal members may have some capacity for lithotrophy in addition to organotrophic growth.

Villa Luz snottites were more diverse than snottites from other caves (Table S1). The abundance of “other bacteria” in Villa Luz (Figure 7, Table 2) indicates that additional diversity in these samples remains to be explored, and the deeper rRNA gene amplicon libraries from this cave contain several taxa that haven’t previously been described from snottites (see also D’Auria et al. (35)). More work is needed to reveal the diversity of extreme acidophiles in this cave community.

### Implications for community assembly

Sulfidic cave snottites from across the globe are populated by extreme acidophiles from the same few genera and families. Within those clades, however, populations are divergent. The *Acidithiobacillus, Sulfobacillus*, and *Acidimicrobiaceae* from snottites in the Mexican caves are less than 95% similar to those from the Italian caves, and *Ferroplasma* spp. are less than 97% similar (Figure 2, 3, 4, S1). Furthermore, based on FISH cell counts, differences among snottite communities are statistically significantly related to cave location (Figure 7), so communities are site specific.

This pattern is consistent with a model of snottite community assembly in which caves are stochastically colonized by microorganisms from local sources, followed by selective filtering for those organisms that can survive in the extremely acidic subaerial cave environment. If colonization events are sufficiently rare, then differences among snottite communities could reflect differences in the founding community and/or later colonizers. Dispersal rates of extreme acidophiles into these caves should be very low: snottites are analogous to extremely acidic “islands” in a “sea” of alkaline limestone bedrock, so survival of extreme acidophiles during transport into the cave environment would be rare. Consistent with this constraint, previous research on the population structure of *Acidithiobacillus* spp. provides strong evidence for restricted dispersal within and among sulfidic caves (36), and studies of other subsurface communities have shown that they retain a site-specific “fingerprint” of their founding communities (e.g., 28). If this model is correct, then differences in the composition of snottite communities are partially the result of historical contingency. Whatever extreme acidophiles happened to enter the cave were those that went on to form and colonize the extremely acidic biofilms. Restricted dispersal could lead to novel diversity in this situation as those populations are free to occupy new niche space and evolve and adapt *in situ*.

Differences among snottite communities could also result from environmental selection. Earlier research found that *Acidithiobacillus* spp. from Frasassi and Acquasanta had differences in their sulfur oxidation pathways and other functional genes (36), which indicates distinct metabolic capabilities that could confer selective advantages in the environment. These two *Acidithiobacillus* populations from Frasassi and Acquasanta had identical 16S rRNA genes. Therefore, the far more divergent *Acidithiobacillus, Ferroplasma, Sulfobacillus*, and *Acidimicrobiaceae* in Mexico versus Italy likely have substantial physiological differences that the cave environment could select for. Snottites were chosen for this study because they occur in a very specific geochemical niche in sulfidic caves (overhanging subaerial cave surfaces, H_2_S(*g*) between 0.2-25 ppm, extremely low pH; Figure 1a), so environmental variability among them should be limited. However, we measured subtle but important geochemical differences: Frasassi was substantially cooler than Acquasanta, Villa Luz, and Luna Azufre, SO_2_(*g*) was present in the atmosphere of Acquasanta but not the other caves at the time of sampling (Figure 7, Table 1), and there are likely other differences in nutrient or energy sources that we did not measure.

Villa Luz snottites were dominated by *Acidithiobacillus* spp. “group 2” that are more closely related to *At. caldus* than *At. thiooxidans*. If *At. thiooxidans* populations were rare but present in Villa Luz snottites, that would be evidence that they were being selected against. However, deep amplicon sequencing did not identify any *At. thiooxidans* out of 246,173 amplicon sequences from Villa Luz snottites, although that does not exclude the possibility that *At. thiooxidans* have had other opportunities to colonize the cave or that they exist elsewhere in the cave. Interestingly, this is not true of the *Acidimicrobiaceae*. The most abundant *Acidimicrobiaceae* OTU from the Villa Luz amplicon libraries clusters with the Villa Luz clones (24.8% and 13.7% of VL13-1 and VL13-2 libraries, respectively), but there is a second rare OTU that is more similar to the Frasassi clones that represents 1.1% and 0.003% of total sequences from these libraries (OTU_9 in Figure 3). Nevertheless, this rare OTU is still >3% different from the Frasassi *Acidimicrobiaceae* clones. Evolutionary rates for 16S rRNA genes are 0.025 to 0.091% per million years (48), so these differences represent at least tens of millions of years of evolution, which highlights how different the Villa Luz and Luna Azufre populations are compared to those in Frasassi and Acquasanta. The formation and colonization of snottites is apparently an adaptation to the sulfidic cave environment that can be exploited by substantially different organisms.

## Methods

### Sample collection and geochemistry

Snottite biofilm samples were collected from four sulfidic caves: Cueva de Villa Luz and Cueva Luna Azufre in Tabasco, Mexico, and the Grotte del Fiume (Frasassi) and Grotta Nuova di Rio Garrafo (Acquasanta Terme) in the Marche region, Italy. Villa Luz is located 2 km south of Tapijulapa in Tabasco, Mexico, and contains roughly 2 km of passage in a Cretaceous limestone. The entrance to Luna Azufre is located 320 meters southeast from Villa Luz, and Luna Azufre has over 500 m of passage in the same formation. Both caves are fed by phreatic water containing up to 15 mM H_2_S(*aq*) that could originate either from nearby volcanic or petroleum sources (30, 49, 50). The Frasassi Caves are located 3 km southeast of Genga in the Marche region, Italy, and contain over 25 km of passage in a Jurassic platform limestone. Sulfidic waters entering Frasassi contain up to 600 μM H_2_S(*aq*) that likely originates from sulfate reduction in an underlying evaporite formation (51, 52). Grotta Nuova di Rio Garrafo is located 2 km south of Acquasanta Terme, and 80 km to the southeast of Frasassi. The Acquasanta cave consists of over 1 km of passage in a marly Eocene limestone, and is fed by thermal sulfidic waters that reach temperatures up to 50°C with over 800 μM H_2_S(*aq*) (37, 53).

Samples were collected from 2-16 locations within each cave. At each sampling site, multiple snottites (approximately 0.5-1.5 grams of biofilm) were collected into sterile centrifuge tubes from an area of cave wall no larger than one square meter. At the time of collection, the pH values of at least 5-10 snottites from the collection area were measured with pH paper (range 0-2.5). The pH of all samples was between 0-1.5.

H_2_S(*g*) concentration was measured at the time of collection using Dräger tubes (range 0.2-60 ppm) (Dräger Safety Inc, Germany) and by an EN-MET MX2100 portable gas detector (range 1-120 ppm) (ENMET corp, USA). SO_2_(*g*) and CO_2_(*g*) concentrations were also measured by ENMET MX2100 (lower detection limits 0.1 ppm and 0.1 %, respectively), and CO_2_(*g*) was also measured by Dräger tubes (100 to 6000 ppm) at some sites. In Villa Luz and Luna Azufre, samples for CO_2_(g) concentration were collected in sealed 60 mL serum bottles (2 bottles per site, flushed 10 times each with 1 volume cave air) and measured with a SRI 310 gas chromatograph (GC) equipped with a thermal conductivity detector and a PoraPak Q packed column (3’ x 1/8”) with helium carrier gas at 80°C (SRI instruments, USA). NH_3_(*g*) and N_2_O(*g*) were measured using Dräger tubes at Acquasanta and Frasassi in 2005-2007, but those gases were never detected (lower detection limit 0.25 and 0.5 ppm, respectively). The total range of multiple gas measurements at each site is given in Table 1.

### 16S rRNA gene libraries

For DNA extraction, approximately 1 gram of snottite biofilm was preserved in 5 parts RNAlater (Ambion, USA) within 4 hours of collection, and stored at - 20°C. DNA extraction of samples RS05-2 and PC05-1 were described in Macalady et al. (2007). For samples RS05-24, VL07-20, LA07-10, AS07-3, AS08-5, and RS09-1, DNA was extracted via bead beating and phenol-chloroform extraction as described in (32, 54). RNAlater-preserved samples were first diluted with 3 parts 1x phosphate-buffered saline (PBS). To remove polysaccharides from the DNA extract, prior to final elution the DNA was resuspended in 200 μL tris (200 mM, pH 8.0), 100 μL NaCl (5 M), and 600 μL ethanol (100%), incubated at -20°C for 30 minutes, and pelleted by centrifugation.

Bacterial, universal, and archaeal 16S rRNA gene clone libraries were constructed from environmental DNA templates. For bacterial and universal libraries, 16S rRNA genes were amplified by polymerase chain reaction (PCR), ligated into pCR4-TOPO vector, and transformed into OneShot Mach1 *E. coli* as described in (31). Archaeal clones were sequenced from sample VL07-20 using forward primer 344f (ACG GGG YGC AGC AGG CGC GA) (55) and reverse primer 1492r (GGT TAC CTT GTT ACG ACT T) (56). Archaeal clones were sequenced from samples RS05-24 and RS09-1 using 344f and reverse primer deg1392r (ACR GGC GGT GTG TRC) (32). Amplification, ligation, and transformation of the archaeal sequences was performed as in Jones et al. (32). Some additional 16S rRNA gene clones were recovered from Frasassi samples GS08-6, GB08-30, and RS08-31 using 27F and primer Sag2 for the *Acidithiobacillus* 16S-23S intergenic transcribed spacer region, as described in (36). Inserts were extracted from RS05-2 and PC05-1 via plasmid extraction as described in (31). Inserts for all other samples were amplified by colony PCR as described in (57). Clones were sequenced at the Penn State University Nucleic Acid Facility.

High-throughput rRNA gene “amplicon” libraries were generated from two snottite samples from Villa Luz collected in 2013 (58). Samples were preserved in sucrose lysis buffer (59), and DNA was extracted using the MoBio PowerSoil™ DNA Isolation Kit (MoBio, Carlsbad, CA). Amplicon libraries were created and sequenced by Molecular Research LP (MR DNA) (Shallowater, TX), using bacterial-specific primers 46f (5′-GCYTAAYACATGCAAGTCG-3′) and 1409r (5′-GTGACGGGCRGTGTGTRCAA-3′) (59, 60).

### Fluorescence *in situ* hybridization (FISH)

Approximately 0.25-0.5 grams of biofilm was fixed for FISH analysis. Samples were either fixed for 3-4 hours in 3% (w/v) paraformaldehyde within 10 hours of collection, or stored in RNAlater (Ambion, USA) as described above and later fixed in paraformaldehyde. After fixation, samples were stored in 1:1 PBS/ethanol at -20°C. Fluorescence *in situ* hybridization followed the procedure outlined by Hugenholtz et al. (61). Environmental samples were applied to 10-well Teflon coated slides, dried, and dehydrated for 3 minutes each in 50%, 80%, and 90% ethanol washes. Hybridization buffer contained 0.9 M NaCl, 20 mM of Tris/HCl pH 7.4, 0.01% sodium dodecyl sulfate (SDS), 25 ng of each fluorescently labeled probe (Table 3), and variable formamide concentrations depending on the probe (Table 3). Slides were incubated at 46°C for 2 hours, followed by a 15 minute incubation at 48°C in wash buffer containing 20 mM Tris/HCl pH 7.4, 0.01% SDS, and variable NaCl concentrations following Lathe (62). Slides were rinsed with deionized water, counterstained with 4’,6’-diamidino-2-phenylindole (DAPI), dried, and mounted with Vectashield (Vector Laboratories Inc, Burlingame, CA, USA). Slides were viewed at 1000x magnification with a Nikon Eclipse 80i epifluorescence microscope, and photomicrographs were taken with a monochrome Photometrics CoolSNAP ES^2^ CCD camera using NIS-Elements AS 3.0 software.

Between 800-1200 DAPI-stained cells were counted for each probe combination. Cells were counted from at least six photos, each representing a different microscope field, from at least two slide wells. Standard deviation was calculated from differences in proportion of probe-labeled cells among each photo. To ensure objectivity in counts, photos for counting were taken without prior examination of the field of view under more than one filter (EUBMIX whenever possible).

#### Phylogenetic, bioinformatic, and statistical analyses

Clone sequences were assembled and edited using CodonCode Aligner v.2.0.4 (CodonCode Corp., USA). Chimeric sequences were identified and removed using Bellerophon 3 (63). Contiguous sequences were aligned using the SINA aligner (64) (https://www.arb-silva.de) and loaded into an ARB (65) database with SILVA database v. 132 (66, 67). Alignments were manually refined using the ARB_Edit4 aligner.

For phylogenetic analysis, alignments were trimmed to equalize sequence length, and positions with <50% gaps were masked. Maximum likelihood phylogenies of full-length sequences were calculated with RAxML v.8.2.12 (68), using the general time reversible nucleotide substitution model, gamma distributed rates, the proportion of invariant sites and base frequencies estimated from the data, and 100 rapid bootstrap replicates. Percent nucleotide sequence similarity was calculated in ARB.

High throughput amplicon libraries were processed using the pipeline described in (69), except that only the first 200 bp of the R1 (forward) read was used for OTU clustering and taxonomic identification. Taxonomic classification of representative OTU sequences was determined with mothur v.1.3.2 using the SILVA database v.132 and a confidence cutoff of 50 (70). Short amplicon sequences were then placed into phylogenetic trees using the evolutionary placement algorithm (EPA) with default parameters (71).

For statistical analyses of FISH cell counts, we used average proportions of THIO1, ACM732, FER656, “other bacteria” (total EUBMIX – (THIO1 + ACM732)), and “other archaea” (ARCH915 – FER656). Principal component analyses (PCA), ANOSIM and Manel tests (72) were performed using the vegan package in R (73, 74). The geographic distance matrix used for Mantel tests included linear distances among individual caves and linear distances between sampling locations within the same cave system, either measured at the time of sampling or from cave maps (Table S3). Mantel statistics were evaluated with Spearman’s rank correlation and 1000 randomizations. For PCA analyses, cross-product matrices were calculated with Pearson’s correlation coefficients.

### Accession number(s)

16S rRNA gene clones referenced in this study are available in the GenBank database under accession numbers KU341124-KU341208, OP269604-OP269649, KC582420-KC582530, and DQ499162-DQ499330. Amplicon libraries are available in NCBI’s Sequence Read Archive under project PRJNA878455.

## Supporting information

Main text

## Acknowledgements

Thanks to A. Montanari for logistical support and the use of facilities and laboratory space at the Osservatorio Geologico di Coldigioco (Italy). S. Mariani, S. Galdenzi, S. Cerioni, and M. Mainiero provided expert advice and field assistance with research in Italy, and F. Baldoni, S. Recanatini, S. Dattagupta, and members of the Gruppo Speleologico CAI di Fabriano and the Gruppo Speleologico Marchigiano CAI di Ancona assisted with sample collection and provided additional excellent field support. We thank L. Rosales-Lagarde, L. Hose, and S. Dattagupta for logistical support, insightful discussion, and field assistance with sample collection in Mexico. Thanks to the municipal government of Tacotalpa, Tabasco, Mexico, and to C. Rogers Morales Mendez and L. Felino Arevalo Gallegos, for granting permits for cave access and sample collection. C. Alberto Cordero Martinez provided lodging and logistical support in Villa Luz Park. The two 2013 Villa Luz samples were collected during an expedition supported by National Geographic grant #EC0644-13 to P. Boston, and sequencing of these two libraries was supported by a Vehslage Grant from the National Speleological Foundation. This work was supported by a graduate research fellowship to DSJ from the Cave Conservancy Foundation, and grants to JLM from the National Science Foundation (EAR 0311854 and EAR 0527046) and NASA NAI (NNA04CC06A).

## Notes

### Competing Interest Statement

The authors have declared no competing interest.

### Summary of Updates

Uploaded accompanying supplementary materials

