## Supplementary material for "Convergent community assembly among globally separated acidic cave biofilms": Main text

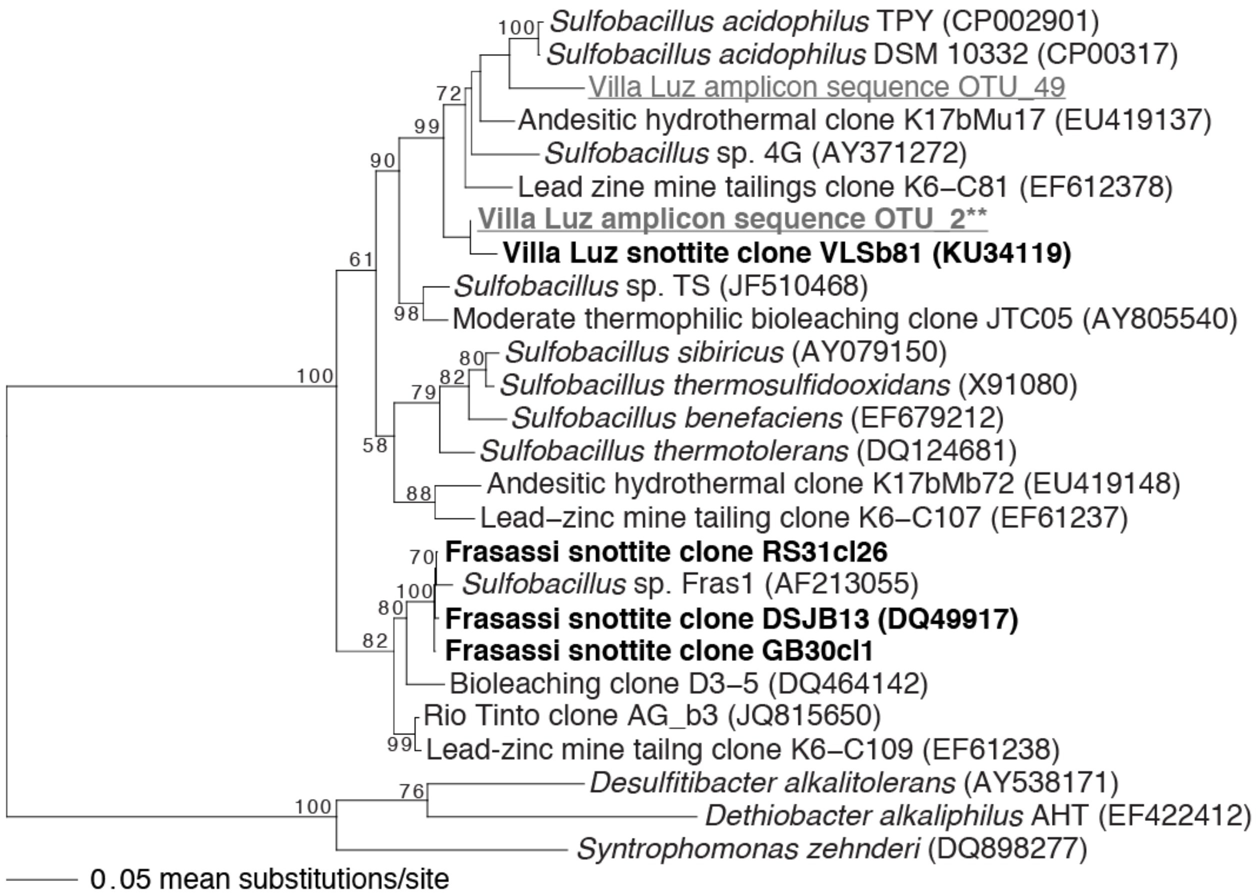


**Figure S1.** Phylogenetic analysis of 16S rRNA gene sequences from genus *Sulfobacillus*. The base tree is a maximum likelihood phylogram created with nearly full-length sequences, with shorter amplicon sequences (gray underline) placed after the fact using the EPA algorithm. Representative sequences from 16S rRNA gene clones are in bold black. The amplicon sequence in bold gray indicated by two asterisks (**) represents that most abundant *Sulfobacillus* OTU (2.2 and 56.1% of the VL13-1 and VL13-2 libraries, respectively); other *Sulfobacillus* OTUs are <0.5% relative abundance (Table S2). Bootstrap values >50% are provided for each node.

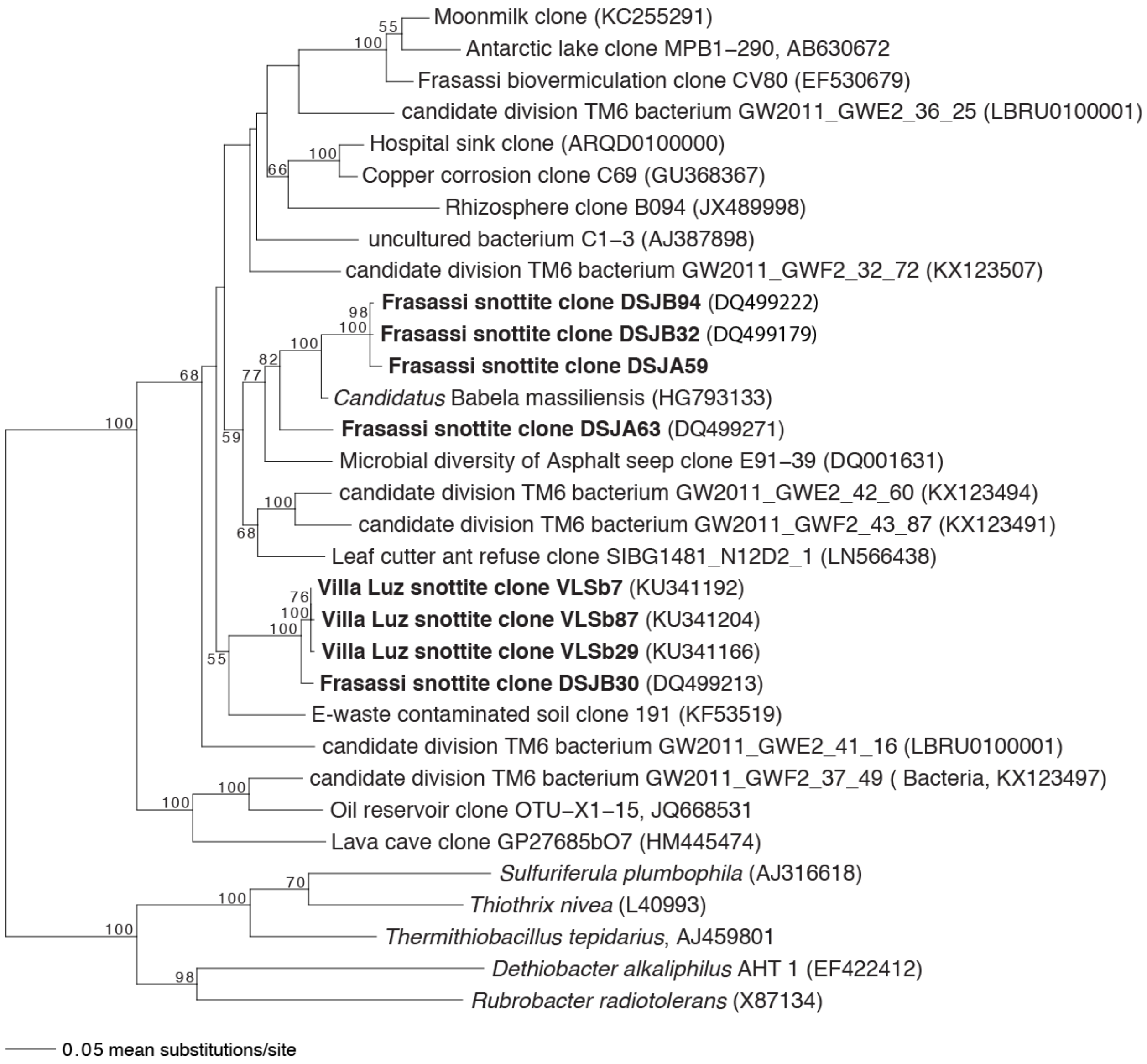


**Figure S2.** Maximum parsimony phylogram of 16S rRNA gene sequences from the *Dependentiae* (formerly TM6 group). Bootstrap values >50% are provided for each node, and snottite sequences from this study are in bold.

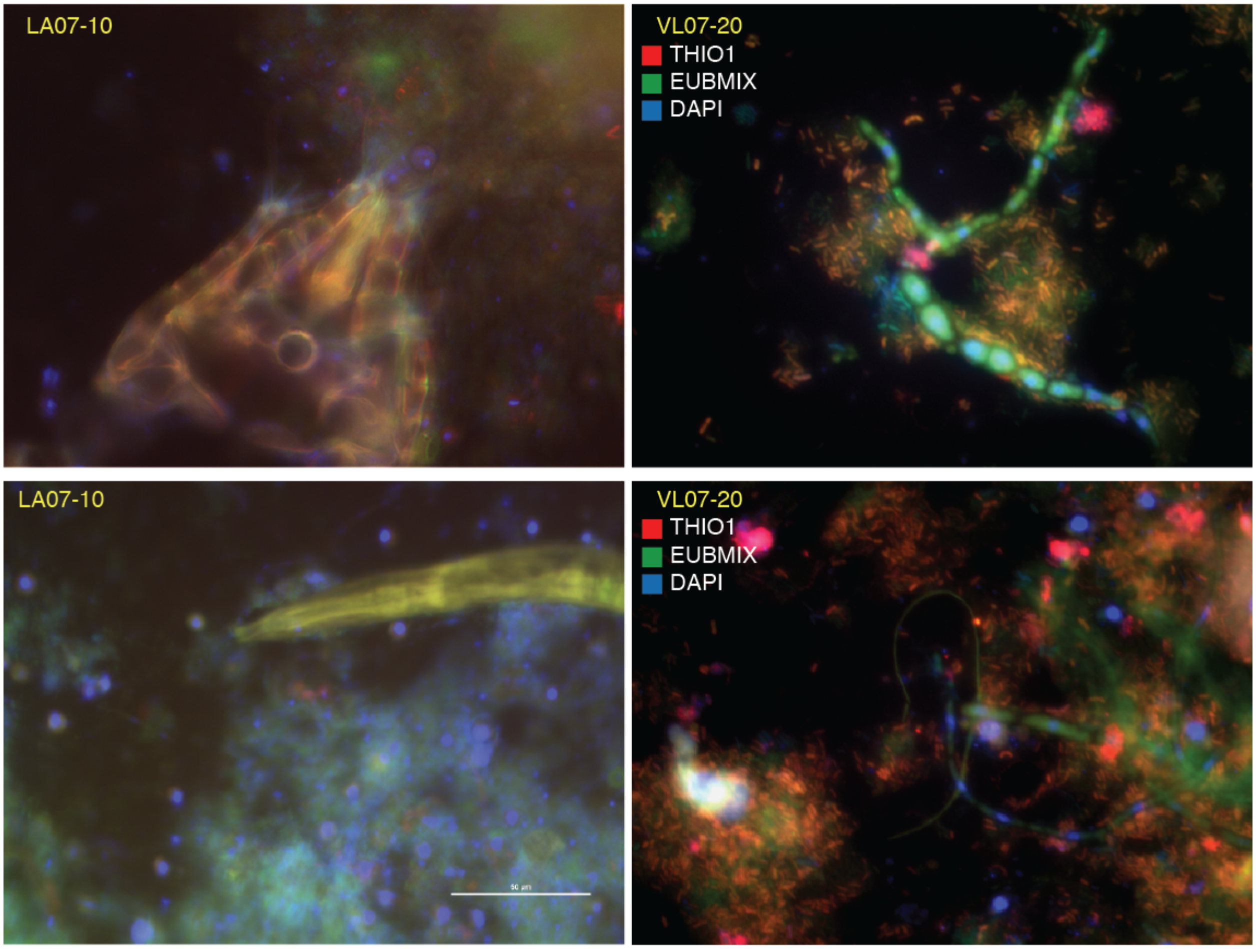


**Figure S3.** Images of eukaryotes observed in Villa Luz and Luna Azufre snottites during epifluorescence microscopy. Upper left image is part of an unknown arthropod; lower left image is part of an unknown nematode; and the large filaments in the right-hand images are likely fungal hyphae.

**Table S1. Summary of 16S rRNA gene clone libraries.**

| Genus | *Acidithiobacillus* | | *Acidimicrobium/ Ferrimicrobium* | | *Sulfobacillus* | | TM6 | | | TM7 | | | Other bacteria[b] | *Ferroplasma* | | C-plasma | D-plasma | G-plasma |
|---|---|---|---|---|---|---|---|---|---|---|---|---|---|---|---|---|---|---|
| Representative clone identifier[a] | DSJb51/ FS43/ ASb9 | LASb10/ VLSb10 | DSBb5 | VLSb84/ LASb14 | DSJb13 | VLSb81 | DSJb94 | DSJa63 | DSJb30/ VLSb7 | VLSb3 | VLSb65 | LASb9 | | DSJa14/ AS5u58 | VLSa11 | DSJa51/ VLSa30 | RS24_ A14 | RS24_ A1 |
| *Bacterial and universal* | | | | | | | | | | | | | | | | | | |
| AS07-3 (ASb clones, n=44) | 44 | | | | | | | | | | | | | | | | | |
| AS08-5 (AS5 clones, n=88) | 70 | | | | | | | | | | | | | 18 | | | | |
| VL07-20 (VLS clones, n=53) | | 20 | | 7 | | 4 | | | 1 | 6 | 2 | | 6 | | 6 | 1 | | |
| LA07-10 (LAS clones, n=47) | | 36 | | 8 | | | | | | | | 2 | 1 | | | | | |
| PC05-1 (FS clones, n=71) | 71 | | | | | | | | | | | | | | | | | |
| RS05-2 (DSJ clones, n=103) | 68 | | 22 | | 1 | | 1 | 1 | 3 | | | | | 6 | | 1 | | |
| *Archaeal-specific* | | | | | | | | | | | | | | | | | | |
| RS05-24 (RS05-24c clones, n=28) | | | | | | | | | | | | | | | | 1 | 1 | 26 |
| RS09-1 (RS9a clones, n=25) | | | | | | | | | | | | | | | | | | 25 |

Numbers refer to the number of clones represented by each phylotype (98% sequence similarity).

[a]This is the name of the clone that presents the phylotype in phylogenetic analyses (Figures 2, 3, 4, S1, and S2).

[b]Other bacteria include phylotypes related to eukaryotic symbionts (VLSb27, VLSb33, VLSb55, VLSb89), Alphaproteobacteria (VLSb58, LASb19), and Firmicutes (VLSb25).

Mitochondrial sequences are not included in this table.

**Table S2.** Relative abundance (%) and taxonomic classification of OTUs from high throughput bacterial rRNA gene amplicon libraries

| OTUId | Sample ID | | Taxonomy | | | | | |
|---|---|---|---|---|---|---|---|---|
| | VL13-1 | VL13-2 | domain | phylum | class | order | family | genus |
| OTU_1 | **48.109** | **24.855** | Bacteria(100) | Proteobacteria(100) | Gammaproteobacteria(100) | Acidithiobacillales(97) | Acidithiobacillaceae(97) | Acidithiobacillus(96) |
| OTU_2 | **2.227** | **56.137** | Bacteria(100) | Firmicutes(100) | Clostridia(100) | Clostridiales(100) | Family_XVII(100) | Sulfobacillus(100) |
| OTU_3 | **24.829** | **13.683** | Bacteria(100) | Actinobacteria(100) | Acidimicrobiia(100) | Acidimicrobiales(100) | Acidimicrobiaceae(100) | uncultured(100) |
| OTU_4 | **16.471** | 0.045 | Bacteria(99) | Proteobacteria(95) | Gammaproteobacteria(93) | Xanthomonadales(77) | Rhodanobacteraceae(76) | Metallibacterium(57) |
| OTU_5 | 0.740 | **4.027** | unknown | unclassified | unclassified | unclassified | unclassified | unclassified |
| OTU_6 | **1.493** | 0.021 | Bacteria(100) | Actinobacteria(99) | Acidimicrobiia(99) | IMCC26256(67) | IMCC26256_fa(67) | IMCC26256_ge(67) |
| OTU_7 | **1.134** | 0.003 | Bacteria(100) | Actinobacteria(100) | Acidimicrobiia(100) | Acidimicrobiales(100) | Acidimicrobiaceae(100) | uncultured(100) |
| OTU_8 | 0.906 | 0.003 | Bacteria(100) | Patescibacteria(100) | Saccharimonadia(100) | Saccharimonadales(100) | Saccharimonadales_fa(100) | Saccharimonadales_ge(100) |
| OTU_9 | 0.703 | 0.039 | Bacteria(95) | Actinobacteria(84) | Acidimicrobiia(72) | unclassified | unclassified | unclassified |
| OTU_10 | 0.317 | 0.017 | Bacteria(100) | Actinobacteria(99) | Actinobacteria(99) | Corynebacteriales(76) | Mycobacteriaceae(62) | Mycobacterium(62) |
| OTU_11 | 0.223 | 0.042 | Bacteria(100) | Bacteroidetes(100) | Bacteroidia(100) | Cytophagales(99) | Amoebophilaceae(99) | Candidatus_Amoebophilus(94) |
| OTU_12 | 0.235 | 0.003 | Bacteria(100) | Patescibacteria(100) | Saccharimonadia(100) | Saccharimonadales(100) | Saccharimonadales_fa(100) | Saccharimonadales_ge(100) |
| OTU_13 | 0.289 | 0.251 | Bacteria(100) | Proteobacteria(100) | Gammaproteobacteria(100) | Vibrionales(100) | Vibrionaceae(100) | Vibrio(99) |
| OTU_14 | 0.269 | 0.019 | Bacteria(100) | Actinobacteria(100) | Thermoleophilia(100) | Solirubrobacterales(97) | Solirubrobacteraceae(92) | Conexibacter(88) |
| OTU_15 | 0.159 | 0.049 | Bacteria(100) | Actinobacteria(99) | Actinobacteria(99) | Corynebacteriales(85) | Mycobacteriaceae(77) | Mycobacterium(77) |
| OTU_16 | 0.150 | 0.015 | Bacteria(98) | Patescibacteria(93) | Saccharimonadia(92) | Saccharimonadales(92) | Saccharimonadales_fa(92) | Saccharimonadales_ge(92) |
| OTU_17 | 0.139 | 0.000 | Bacteria(100) | Actinobacteria(100) | Actinobacteria(100) | Corynebacteriales(100) | Mycobacteriaceae(100) | Mycobacterium(100) |
| OTU_18 | 0.103 | 0.000 | Bacteria(100) | Proteobacteria(100) | Alphaproteobacteria(100) | Acetobacterales(100) | Acetobacteraceae(100) | Acidiphilium(77) |
| OTU_19 | 0.089 | 0.008 | Bacteria(100) | Proteobacteria(100) | Alphaproteobacteria(100) | Paracaedibacterales(100) | Paracaedibacteraceae(100) | uncultured(100) |
| OTU_20 | 0.050 | 0.064 | Bacteria(100) | Firmicutes(100) | Clostridia(100) | Clostridiales(100) | Clostridiaceae_1(100) | Clostridium_sensu_stricto_1(98) |
| OTU_21 | 0.062 | 0.055 | Bacteria(100) | Proteobacteria(100) | Gammaproteobacteria(100) | Pseudomonadales(100) | Pseudomonadaceae(100) | Pseudomonas(100) |
| OTU_22 | 0.035 | 0.040 | Bacteria(100) | Firmicutes(100) | Clostridia(100) | Clostridiales(100) | Clostridiaceae_1(100) | Clostridium_sensu_stricto_1(98) |
| OTU_23 | 0.119 | 0.001 | Bacteria(100) | Actinobacteria(100) | Acidimicrobiia(100) | uncultured(100) | uncultured_fa(100) | uncultured_ge(100) |
| OTU_24 | 0.065 | 0.000 | Bacteria(99) | Actinobacteria(92) | Acidimicrobiia(87) | Acidimicrobiales(87) | Acidimicrobiaceae(87) | unclassified |
| OTU_25 | 0.057 | 0.000 | Bacteria(100) | Patescibacteria(100) | Saccharimonadia(100) | Saccharimonadales(100) | Saccharimonadales_fa(100) | Saccharimonadales_ge(100) |
| OTU_26 | 0.030 | 0.019 | Bacteria(100) | Proteobacteria(100) | Gammaproteobacteria(100) | Aeromonadales(100) | Aeromonadaceae(100) | Aeromonas(99) |
| OTU_27 | 0.044 | 0.019 | Bacteria(100) | Proteobacteria(100) | Gammaproteobacteria(100) | Aeromonadales(100) | Aeromonadaceae(100) | Aeromonas(100) |
| OTU_28 | 0.016 | 0.030 | Bacteria(100) | Actinobacteria(100) | Acidimicrobiia(100) | uncultured(76) | uncultured_fa(76) | uncultured_ge(76) |
| OTU_29 | 0.021 | 0.016 | Bacteria(100) | Proteobacteria(100) | Gammaproteobacteria(100) | Betaproteobacteriales(100) | Rhodocyclaceae(87) | uncultured(80) |
| OTU_30 | 0.017 | 0.023 | Bacteria(97) | Acidobacteria(83) | Acidobacteriia(77) | Subgroup_2(77) | Subgroup_2_fa(77) | Subgroup_2_ge(77) |
| OTU_31 | 0.033 | 0.009 | Bacteria(100) | Proteobacteria(100) | Gammaproteobacteria(100) | KF-JG30-C25(93) | KF-JG30-C25_fa(93) | KF-JG30-C25_ge(93) |
| OTU_32 | 0.043 | 0.038 | Bacteria(100) | Proteobacteria(100) | Gammaproteobacteria(100) | Enterobacteriales(100) | Enterobacteriaceae(100) | unclassified |
| OTU_33 | 0.027 | 0.006 | Bacteria(95) | Actinobacteria(88) | Acidimicrobiia(81) | unclassified | unclassified | unclassified |
| OTU_34 | 0.022 | 0.015 | Bacteria(100) | Proteobacteria(100) | Gammaproteobacteria(100) | Acidiferrobacterales(100) | Acidiferrobacteraceae(100) | Sulfurifustis(100) |
| OTU_35 | 0.040 | 0.000 | Bacteria(97) | Actinobacteria(63) | Acidimicrobiia(60) | unclassified | unclassified | unclassified |
| OTU_36 | 0.038 | 0.018 | Bacteria(100) | Actinobacteria(100) | Actinobacteria(100) | Corynebacteriales(100) | Mycobacteriaceae(84) | Mycobacterium(84) |
| OTU_37 | 0.017 | 0.014 | Bacteria(100) | Proteobacteria(100) | Alphaproteobacteria(100) | Rhodobacterales(100) | Rhodobacteraceae(100) | Thioclava(98) |
| OTU_38 | 0.024 | 0.018 | Bacteria(100) | Epsilonbacteraeota(100) | Campylobacteria(100) | Campylobacterales(100) | Sulfurovaceae(86) | Sulfurovum(82) |
| OTU_39 | 0.022 | 0.022 | Bacteria(100) | Proteobacteria(100) | Gammaproteobacteria(100) | Alteromonadales(100) | Shewanellaceae(100) | Shewanella(100) |
| OTU_40 | 0.006 | 0.009 | Bacteria(81) | unclassified | unclassified | unclassified | unclassified | unclassified |
| OTU_41 | 0.015 | 0.000 | Bacteria(100) | Proteobacteria(100) | Gammaproteobacteria(100) | Legionellales(100) | Legionellaceae(100) | Legionella(100) |
| OTU_42 | 0.013 | 0.011 | Bacteria(100) | Actinobacteria(100) | Actinobacteria(100) | Corynebacteriales(91) | Mycobacteriaceae(81) | Mycobacterium(81) |
| OTU_43 | 0.016 | 0.000 | Bacteria(100) | Dependentiae(100) | Babeliae(100) | Babeliales(100) | Babeliaceae(93) | Babeliaceae_ge(85) |
| OTU_44 | 0.067 | 0.036 | Bacteria(100) | Proteobacteria(100) | Gammaproteobacteria(100) | Enterobacteriales(100) | Enterobacteriaceae(100) | Lelliottia(85) |
| OTU_45 | 0.009 | 0.013 | Bacteria(87) | Firmicutes(65) | Bacilli(52) | Bacillales(51) | unclassified | unclassified |
| OTU_46 | 0.015 | 0.000 | Bacteria(72) | unclassified | unclassified | unclassified | unclassified | unclassified |
| OTU_47 | 0.022 | 0.029 | Bacteria(100) | Proteobacteria(100) | Alphaproteobacteria(100) | Rhodospirillales(100) | Thalassospiraceae(100) | Thalassospira(100) |
| OTU_48 | 0.005 | 0.005 | Bacteria(93) | Bacteroidetes(88) | Bacteroidia(88) | Cytophagales(78) | Microscillaceae(64) | uncultured(56) |
| OTU_49 | 0.018 | 0.001 | Bacteria(97) | Firmicutes(94) | Clostridia(94) | Clostridiales(94) | Family_XVII(94) | Sulfobacillus(94) |
| OTU_50 | 0.011 | 0.004 | Bacteria(100) | Proteobacteria(100) | Gammaproteobacteria(100) | Betaproteobacteriales(100) | Hydrogenophilaceae(100) | Thiobacillus(100) |
| OTU_51 | 0.022 | 0.000 | Bacteria(100) | Proteobacteria(100) | Gammaproteobacteria(100) | Gammaproteobacteria_Incertae_Sedis(98) | Unknown_Family(98) | Acidibacter(98) |
| OTU_52 | 0.005 | 0.003 | Bacteria(93) | Chloroflexi(86) | Dehalococcoidia(86) | GIF9(81) | AB-539-J10(74) | AB-539-J10_ge(73) |
| OTU_53 | 0.000 | 0.011 | Bacteria(100) | Bacteroidetes(100) | Bacteroidia(100) | Cytophagales(97) | Amoebophilaceae(97) | Candidatus_Amoebophilus(69) |
| OTU_54 | 0.012 | 0.000 | Bacteria(100) | Dependentiae(100) | Babeliae(100) | Babeliales(100) | UBA12409(100) | UBA12409_ge(100) |
| OTU_55 | 0.022 | 0.008 | Bacteria(100) | Proteobacteria(100) | Gammaproteobacteria(100) | Betaproteobacteriales(100) | Burkholderiaceae(95) | Thiomonas(78) |
| OTU_56 | 0.039 | 0.015 | Bacteria(100) | Proteobacteria(100) | Gammaproteobacteria(100) | Alteromonadales(96) | Shewanellaceae(96) | Shewanella(96) |
| OTU_57 | 0.015 | 0.008 | Bacteria(100) | Proteobacteria(99) | Gammaproteobacteria(98) | unclassified | unclassified | unclassified |
| OTU_58 | 0.010 | 0.010 | Bacteria(100) | Firmicutes(100) | Bacilli(100) | Bacillales(100) | Bacillaceae(100) | Bacillus(80) |
| OTU_59 | 0.007 | 0.009 | Bacteria(100) | Proteobacteria(97) | Deltaproteobacteria(96) | Desulfobacterales(94) | Desulfobulbaceae(93) | Desulfocapsa(86) |
| OTU_60 | 0.000 | 0.006 | Bacteria(77) | Proteobacteria(70) | Alphaproteobacteria(69) | Rickettsiales(69) | Mitochondria(68) | Mitochondria_ge(68) |
| OTU_61 | 0.010 | 0.006 | Bacteria(100) | Proteobacteria(100) | Gammaproteobacteria(100) | Betaproteobacteriales(100) | Burkholderiaceae(100) | Thiomonas(100) |

| | | | | | | | | |
|---|---|---|---|---|---|---|---|---|
| OTU_62 | 0.008 | 0.006 | Bacteria(100) | Proteobacteria(100) | Gammaproteobacteria(100) | Enterobacteriales(100) | Enterobacteriaceae(100) | unclassified |
| OTU_63 | 0.007 | 0.001 | Bacteria(96) | Actinobacteria(84) | Acidimicrobiia(81) | Acidimicrobiia_or(62) | Acidimicrobiia_fa(62) | Acidimicrobiia_ge(62) |
| OTU_64 | 0.005 | 0.001 | Bacteria(94) | Epsilonbacteraeota(86) | Campylobacteria(86) | Campylobacterales(86) | unclassified | unclassified |
| OTU_65 | 0.006 | 0.006 | Bacteria(76) | unclassified | unclassified | unclassified | unclassified | unclassified |
| OTU_66 | 0.005 | 0.000 | Bacteria(100) | Dependentiae(100) | Babeliae(100) | Babeliales(100) | UBA12409(89) | UBA12409_ge(89) |
| OTU_67 | 0.004 | 0.010 | Bacteria(100) | Actinobacteria(100) | Actinobacteria(100) | Corynebacteriales(100) | Mycobacteriaceae(71) | Mycobacterium(71) |
| OTU_68 | 0.002 | 0.004 | Bacteria(97) | Proteobacteria(93) | Deltaproteobacteria(92) | Desulfobacterales(89) | Desulfobulbaceae(89) | Dissulfurimicrobium(74) |
| OTU_69 | 0.058 | 0.000 | Bacteria(78) | unclassified | unclassified | unclassified | unclassified | unclassified |
| OTU_70 | 0.005 | 0.003 | Bacteria(100) | Proteobacteria(97) | Deltaproteobacteria(97) | Desulfuromonadales(96) | Geobacteraceae(95) | Geobacter(95) |
| OTU_71 | 0.003 | 0.001 | Bacteria(100) | Proteobacteria(100) | Gammaproteobacteria(100) | Xanthomonadales(100) | Rhodanobacteraceae(100) | Chujaibacter(100) |
| OTU_72 | 0.000 | 0.003 | Eukaryota(76) | unclassified | unclassified | unclassified | unclassified | unclassified |
| OTU_73 | 0.000 | 0.006 | Bacteria(88) | TA06(60) | TA06_cl(60) | TA06_or(60) | TA06_fa(60) | TA06_ge(60) |
| OTU_74 | 0.002 | 0.003 | Bacteria(100) | Proteobacteria(100) | Alphaproteobacteria(100) | Micropepsales(92) | Micropepsaceae(92) | uncultured(85) |
| OTU_75 | 0.007 | 0.007 | Bacteria(100) | Firmicutes(100) | Clostridia(100) | Clostridiales(100) | Clostridiaceae_1(100) | Clostridium_sensu_stricto_13(99) |
| OTU_76 | 0.013 | 0.010 | Bacteria(100) | Proteobacteria(100) | Gammaproteobacteria(100) | Enterobacteriales(100) | Enterobacteriaceae(100) | Citrobacter(65) |
| OTU_77 | 0.000 | 0.003 | Bacteria(100) | Chloroflexi(99) | OLB14(99) | OLB14_or(99) | OLB14_fa(99) | OLB14_ge(99) |
| OTU_78 | 0.002 | 0.004 | Bacteria(100) | Chloroflexi(99) | Ktedonobacteria(99) | Ktedonobacterales(96) | Ktedonobacteraceae(96) | HSB_OF53-F07(95) |
| OTU_79 | 0.002 | 0.003 | Bacteria(100) | Firmicutes(100) | Clostridia(100) | Clostridiales(100) | Family_XII(100) | Fusibacter(96) |
| OTU_80 | 0.002 | 0.007 | Bacteria(100) | Bacteroidetes(100) | Bacteroidia(100) | Flavobacteriales(100) | Flavobacteriaceae(100) | Flavobacterium(74) |
| OTU_81 | 0.007 | 0.004 | Bacteria(100) | Proteobacteria(100) | Gammaproteobacteria(100) | Enterobacteriales(100) | Enterobacteriaceae(100) | Erwinia(76) |
| OTU_82 | 0.001 | 0.003 | Bacteria(100) | Firmicutes(100) | Bacilli(99) | Bacillales(99) | Bacillaceae(94) | Bacillus(82) |
| OTU_83 | 0.004 | 0.000 | Bacteria(98) | Proteobacteria(95) | Gammaproteobacteria(94) | Diplorickettsiales(93) | Diplorickettsiaceae(93) | Aquicella(92) |
| OTU_84 | 0.005 | 0.001 | Bacteria(67) | Proteobacteria(65) | Alphaproteobacteria(54) | Rickettsiales(54) | Mitochondria(53) | Mitochondria_ge(53) |
| OTU_85 | 0.003 | 0.001 | Bacteria(100) | Proteobacteria(100) | Gammaproteobacteria(100) | unclassified | unclassified | unclassified |
| OTU_86 | 0.001 | 0.002 | Bacteria(100) | Acidobacteria(100) | Acidobacteriia(100) | Acidobacteriales(100) | Acidobacteriaceae_(Subgroup_1)(100) | Occallatibacter(97) |
| OTU_87 | 0.002 | 0.005 | Bacteria(100) | Proteobacteria(100) | Gammaproteobacteria(100) | Vibrionales(100) | Vibrionaceae(100) | Vibrio(85) |
| OTU_88 | 0.006 | 0.001 | Bacteria(100) | Proteobacteria(100) | Alphaproteobacteria(100) | Caedibacterales(100) | Caedibacteraceae(100) | Caedibacter(100) |
| OTU_89 | 0.011 | 0.001 | Bacteria(100) | Proteobacteria(100) | Gammaproteobacteria(100) | Xanthomonadales(100) | Rhodanobacteraceae(100) | Oleiagrimonas(100) |
| OTU_90 | 0.000 | 0.001 | Bacteria(99) | Bacteroidetes(99) | Bacteroidia(99) | Cytophagales(99) | Microscillaceae(99) | uncultured(98) |
| OTU_91 | 0.003 | 0.002 | Bacteria(83) | Chloroflexi(56) | unclassified | unclassified | unclassified | unclassified |
| OTU_92 | 0.001 | 0.001 | Bacteria(65) | unclassified | unclassified | unclassified | unclassified | unclassified |
| OTU_93 | 0.003 | 0.003 | Bacteria(100) | Firmicutes(100) | Bacilli(100) | Bacillales(100) | Bacillaceae(100) | Fictibacillus(100) |
| OTU_94 | 0.003 | 0.003 | Bacteria(100) | Proteobacteria(100) | Gammaproteobacteria(100) | Acidiferrobacterales(100) | Acidiferrobacteraceae(100) | Sulfurifustis(100) |
| OTU_95 | 0.002 | 0.001 | Bacteria(100) | Firmicutes(100) | Bacilli(100) | Bacillales(100) | Bacillaceae(100) | Fictibacillus(100) |
| OTU_96 | 0.001 | 0.002 | Bacteria(96) | Proteobacteria(86) | Deltaproteobacteria(79) | Desulfobacterales(74) | Desulfobulbaceae(74) | unclassified |
| OTU_97 | 0.000 | 0.001 | Bacteria(100) | Proteobacteria(100) | Gammaproteobacteria(100) | Alteromonadales(99) | Shewanellaceae(95) | Shewanella(95) |
| OTU_98 | 0.003 | 0.003 | Bacteria(98) | Chloroflexi(92) | Anaerolineae(91) | RBG-13-54-9(63) | RBG-13-54-9_fa(63) | RBG-13-54-9_ge(63) |
| OTU_99 | 0.004 | 0.001 | Bacteria(98) | Bacteroidetes(98) | Bacteroidia(98) | Bacteroidales(75) | unclassified | unclassified |
| OTU_100 | 0.002 | 0.001 | Bacteria(100) | Proteobacteria(97) | Gammaproteobacteria(94) | unclassified | unclassified | unclassified |
| OTU_101 | 0.001 | 0.003 | Bacteria(100) | Epsilonbacteraeota(100) | Campylobacteria(100) | Campylobacterales(100) | Arcobacteraceae(100) | Arcobacter(100) |
| OTU_102 | 0.000 | 0.002 | Bacteria(100) | Bacteroidetes(100) | Ignavibacteria(100) | Ignavibacteriales(100) | Ignavibacteriaceae(99) | Ignavibacterium(99) |
| OTU_103 | 0.002 | 0.001 | Bacteria(100) | WS1(100) | WS1_cl(100) | WS1_or(100) | WS1_fa(100) | WS1_ge(100) |
| OTU_104 | 0.002 | 0.001 | Bacteria(96) | Bacteroidetes(94) | Bacteroidia(94) | Cytophagales(76) | Cyclobacteriaceae(67) | unclassified |
| OTU_105 | 0.001 | 0.001 | Bacteria(100) | Acidobacteria(100) | Blastocatellia_(Subgroup_4)(100) | 11-24(100) | 11-24_fa(100) | 11-24_ge(100) |
| OTU_106 | 0.001 | 0.003 | Bacteria(100) | Proteobacteria(100) | Alphaproteobacteria(100) | Elsterales(98) | uncultured(98) | uncultured_ge(98) |
| OTU_107 | 0.005 | 0.001 | Bacteria(98) | Proteobacteria(98) | Alphaproteobacteria(97) | Dongiales(96) | Dongiaceae(96) | Dongia(96) |
| OTU_108 | 0.003 | 0.000 | Bacteria(96) | Actinobacteria(74) | Acidimicrobiia(71) | Acidimicrobiales(50) | unclassified | unclassified |
| OTU_109 | 0.002 | 0.002 | Bacteria(100) | Nitrospirae(100) | Nitrospira(100) | Nitrospirales(100) | Nitrospiraceae(100) | Nitrospira(100) |
| OTU_110 | 0.003 | 0.000 | Bacteria(96) | Proteobacteria(88) | Gammaproteobacteria(78) | Gammaproteobacteria_Incertae_Sedis(67) | Unknown_Family(67) | Candidatus_Ovatusbacter(67) |
| OTU_111 | 0.036 | 0.026 | Bacteria(100) | Proteobacteria(100) | Gammaproteobacteria(100) | Alteromonadales(100) | Shewanellaceae(100) | Shewanella(100) |
| OTU_112 | 0.004 | 0.000 | Bacteria(89) | Proteobacteria(61) | Gammaproteobacteria(57) | unclassified | unclassified | unclassified |
| OTU_113 | 0.004 | 0.003 | Bacteria(100) | Epsilonbacteraeota(97) | Campylobacteria(97) | Campylobacterales(97) | Sulfurovaceae(68) | Sulfurovum(55) |
| OTU_114 | 0.001 | 0.001 | Bacteria(100) | Proteobacteria(100) | Gammaproteobacteria(100) | Betaproteobacteriales(100) | Burkholderiaceae(100) | Castellaniella(100) |
| OTU_115 | 0.006 | 0.006 | Bacteria(99) | Proteobacteria(92) | Deltaproteobacteria(92) | Desulfobacterales(89) | Desulfobulbaceae(87) | Desulfobulbus(58) |
| OTU_116 | 0.004 | 0.002 | Bacteria(100) | Proteobacteria(100) | Alphaproteobacteria(100) | Acetobacterales(100) | Acetobacteraceae(100) | Acidicaldus(62) |
| OTU_117 | 0.001 | 0.001 | Bacteria(100) | Proteobacteria(100) | Gammaproteobacteria(100) | Betaproteobacteriales(100) | SC-I-84(96) | SC-I-84_ge(96) |
| OTU_118 | 0.001 | 0.001 | Bacteria(81) | unclassified | unclassified | unclassified | unclassified | unclassified |
| OTU_119 | 0.004 | 0.003 | Bacteria(100) | Firmicutes(100) | Bacilli(100) | Bacillales(100) | Family_XII(100) | Exiguobacterium(100) |
| OTU_120 | 0.003 | 0.001 | Bacteria(100) | Proteobacteria(100) | Gammaproteobacteria(100) | Betaproteobacteriales(100) | Burkholderiaceae(100) | Leptothrix(83) |
| OTU_121 | 0.003 | 0.001 | Bacteria(98) | Chloroflexi(84) | Anaerolineae(84) | unclassified | unclassified | unclassified |
| OTU_122 | 0.002 | 0.002 | Bacteria(100) | Proteobacteria(100) | Gammaproteobacteria(100) | Betaproteobacteriales(99) | unclassified | unclassified |
| OTU_123 | 0.002 | 0.002 | Bacteria(98) | Epsilonbacteraeota(75) | Campylobacteria(75) | Campylobacterales(73) | unclassified | unclassified |
| OTU_124 | 0.002 | 0.001 | Bacteria(100) | Proteobacteria(100) | Deltaproteobacteria(100) | Sva0485(100) | Sva0485_fa(100) | Sva0485_ge(100) |
| OTU_125 | 0.004 | 0.000 | Bacteria(100) | Proteobacteria(100) | Gammaproteobacteria(100) | unclassified | unclassified | unclassified |
| OTU_126 | 0.001 | 0.003 | Bacteria(98) | Proteobacteria(97) | Gammaproteobacteria(97) | Halothiobacillales(94) | Halothiobacillaceae(94) | Halothiobacillus(94) |
| OTU_127 | 0.003 | 0.000 | Bacteria(100) | Nitrospirae(95) | Nitrospira(95) | Nitrospirales(95) | Nitrospiraceae(95) | Nitrospira(95) |
| OTU_128 | 0.003 | 0.001 | Bacteria(100) | Firmicutes(100) | Bacilli(100) | Bacillales(100) | Bacillaceae(100) | Fictibacillus(100) |
| OTU_129 | 0.005 | 0.001 | Bacteria(95) | Proteobacteria(94) | Gammaproteobacteria(89) | unclassified | unclassified | unclassified |

OTUs with relative abundance >1% displayed in bold font

**Table S3.** Geographic distance separating sampling locations

| | *GS2* | *GS1* | *RS1* | *RS2* | *RS3* | *PC1* | *GB1* | *AS4* | *AS2* | *AS3* | *VL1* | *VL2* | *LA1* | *LA2* |
|---|---|---|---|---|---|---|---|---|---|---|---|---|---|---|
| *GS2* | 0 | | | | | | | | | | | | | |
| *GS1* | 10 m | 0 | | | | | | | | | | | | |
| *RS1* | 215 m | 215 m | 0 | | | | | | | | | | | |
| *RS2* | 215 m | 215 m | 10 m | 0 | | | | | | | | | | |
| *RS3* | 215 m | 215 m | 10 m | 3 m | 0 | | | | | | | | | |
| *PC1* | 150 m | 150 m | 140 m | 140 m | 140 m | 0 | | | | | | | | |
| *GB1* | 50 m | 50 m | 175 m | 175 m | 175 m | 100 m | 0 | | | | | | | |
| *AS4* | 81 km | 81 km | 81 km | 81 km | 81 km | 81 km | 81 km | 0 | | | | | | |
| *AS2* | 81 km | 81 km | 81 km | 81 km | 81 km | 81 km | 81 km | 2 m | 0 | | | | | |
| *AS3* | 81 km | 81 km | 81 km | 81 km | 81 km | 81 km | 81 km | 11 m | 12 m | 0 | | | | |
| *VL1* | 9897 km | 9897 km | 9897 km | 9897 km | 9897 km | 9897 km | 9897 km | 9959 km | 9959 km | 9959 km | 0 | | | |
| *VL2* | 9897 km | 9897 km | 9897 km | 9897 km | 9897 km | 9897 km | 9897 km | 9959 km | 9959 km | 9959 km | 25 m | 0 | | |
| *LA1* | 9897 km | 9897 km | 9897 km | 9897 km | 9897 km | 9897 km | 9897 km | 9959 km | 9959 km | 9959 km | 320 m | 320 m | 0 | |
| *LA2* | 9897 km | 9897 km | 9897 km | 9897 km | 9897 km | 9897 km | 9897 km | 9959 km | 9959 km | 9959 km | 320 m | 320 m | 8 m | 0 |
